# Conditional fusogenic lipid nanocarriers for cytosolic delivery of macromolecular therapeutics

**DOI:** 10.1101/2024.12.27.630514

**Authors:** Qian Zhong, Edward K.W. Tan, Ananth Shyamal, Chayanon Ngambenjawong, Tiziana Parisi, Henry Ko, Heather E. Fleming, Tyler Jacks, Sangeeta N. Bhatia

## Abstract

Macromolecular therapeutics designed for intracellular targets must overcome systemic delivery barriers, target cell membrane impermeability, and inefficient endosomal escape. Here, we engineer a class of conditional fusogenic liposomes (C-FLIPs) that harness the catalytic activity of extracellular proteases present in the pathological microenvironment. This context-specific activation enables on-target membrane fusion with cells in diseased tissue, resulting in cytosolic delivery of therapeutic payloads. We describe the cytoplasmic delivery of three prototypic macromolecular therapeutic classes: peptide degraders, cytotoxic proteins, and ribonucleoprotein particles (RNPs). We further develop C-FLIP to deliver granzyme B (GzmB) to the cytoplasm of cancer cells *in vivo* and induce pyroptosis in immunologically-inert tumors. Treatment with C-FLIP/GzmB reprograms the immunosuppressive tumor microenvironment and synergizes with checkpoint blockade to result in the regression of established tumors and induce immunological memory. This modular, non-viral, cytosolic delivery platform represents a promising approach to leverage pathological protease activity for targeted delivery of biologics.

## Main

The delivery of macromolecular therapeutics to the intracellular environment could enable clinical applications in oncology, as well as in metabolic and hematologic diseases.^1^ However, many biomolecular therapeutics are unable to cross cellular membranes, rendering genetically-validated targets nearly “undruggable”. Propensity for undesirable off-target delivery limits the achievable therapeutic index, even in cases where preclinical candidates demonstrate robust efficacy at the cellular level. It is thus critical to address both physiological and cellular barriers for the delivery of these biomolecular therapeutics. Previously, polymeric and lipid-based nanocarriers have been designed to transport biologics across the cell membrane.^2–5^ Once inside, most nanocarriers are predominantly trapped in endosomal compartments where they are degraded or recycled, with only ∼2% of the payload released into the cytoplasm.^6,7^ Furthermore, nanocarriers including lipid nanoparticles (LNPs) are most effective in delivering anionic payloads.^8,9^ The encapsulation of proteins such as positively-charged Cas9 has proven especially challenging. A universal nanocarrier system for the efficient cytosolic delivery of hydrophilic payloads could accelerate the development of new therapeutic modalities for patients.

To overcome the endosomal barrier, the majority of existing nanocarriers are designed to induce membrane destabilization or osmotic pressure change and thus enable ‘endosomal escape’. However, this mechanism of cytoplasmic delivery is inefficient and accompanied by inherent concomitant toxicity associated with the release of endolysosomal constituents. Alternative strategies include amphiphilic vehicles^10,11^ or cell-penetrating peptides (CPPs)^12,13^ that facilitate direct membrane translocation, yet these methods still face limited delivery efficiency, poor scalability, and weak safety profiles. Inspired by pathogens that bypass receptor-mediated endocytic pathways via direct fusion to deliver genetic codes into the cytoplasm of host cells,^14–16^ lipids have been formulated to enable liposome fusion with cell membranes, promoting a more rapid and efficient delivery of therapeutic payloads. Such fusogenic liposomes (FLIPs) have been used to transfer proteins,^17^ RNA,^18,19^ and subcellular organelles^20^ into target cells, or to enable cell membrane modifications.^21,22^ However, FLIPs indiscriminately fuse with any nearby cells, limiting both efficacy in the diseased tissue and systemic safety.

In this study, we harnessed the catalytic activity of disease-associated proteases to engineer conditional FLIPs (C-FLIPs) for local cytosolic delivery of a range of macromolecules. The C-FLIP is a four-component nanosystem that comprises (i) fusion masks to prevent membrane fusion, (ii) protease substrate tethers designed for conditional cleavage to release fusion masks at diseased sites (termed ‘activation’), (iii) fusogenic liposomes that fuse with cell membranes upon activation, and (iv) encapsulated therapeutic biologics. Upon systemic delivery, we envision that C-FLIPs are activated in diseased tissue resulting in membrane fusion and delivery of the macromolecular payloads into the cytoplasm of target cells, whereas in healthy tissue inactive C-FLIPs are either prevented from entry, or internalized in their inactive form and subsequently degraded (**Fig.1**). C-FLIP payloads can range in size from small molecules or peptides to protein therapeutics and RNPs, as long as they can be formulated in a liposome. Inspired by the cytotoxicity of CD8+ T-cells, we applied C-FLIP to explore the therapeutic potential of selective, cytoplasmic delivery of the cationic cytotoxic enzyme, granzyme B (C-FLIP/GzmB). We found that the GzmB-bearing C-FLIPs induced pyroptosis, an immunogenic form of cell death, in tumors in a protease-dependent fashion. This approach significantly enhanced checkpoint inhibition in an immunologically ‘cold’ mouse model of lung cancer. C-FLIP-mediated delivery, alongside emerging maps of protease dysregulation in disease,^23^ offers a promising non-viral, modular approach for next-generation medicines.

**Figure 1.**
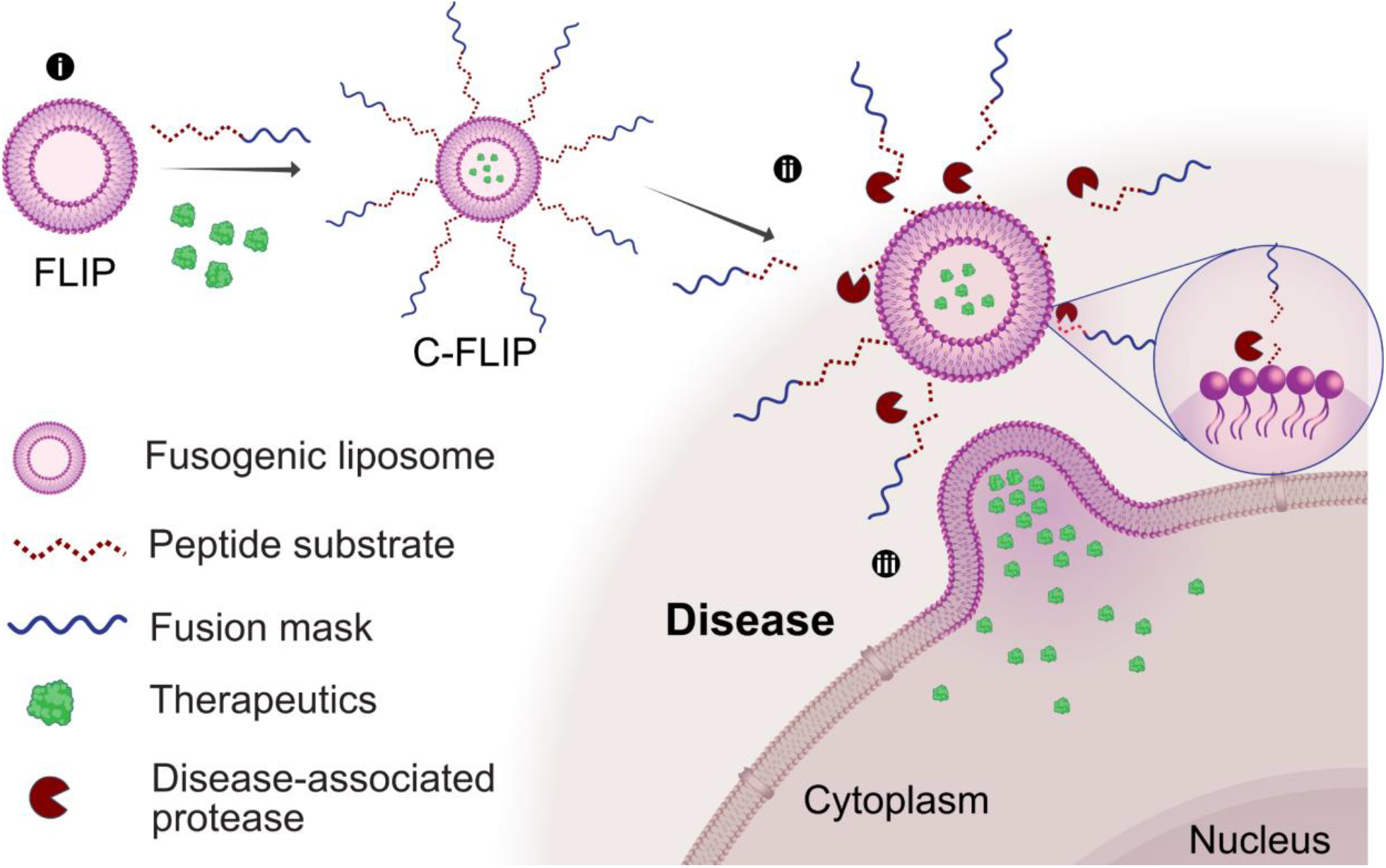
Protease-activated, conditional fusogenic liposomes enable selective, intracellular macromolecular therapeutic delivery. **(i)** Fusogenic liposomes (FLIPs), with encapsulated therapeutics (green), are conjugated to fusion masks via protease-cleavable peptide substrates to form conditional FLIPs (C-FLIPs), **(ii)** Upon systemic delivery, C-FLIP extravasates and disease-specific proteases cleave the substrate linkers, remove the fusion masks, and **(iii)** trigger liposome fusion with the nearby cell membranes to achieve cytosolic delivery of the therapeutics in the pathological microenvironment.

### C-FLIPs exhibit protease-dependent membrane fusion

Threshold levels of lipid fluidity combined with modest cationic charge of liposomes are key factors in promoting membrane fusion when they are proximal to negatively-charged cells.^18^ On this basis, our design of FLIP nanocarriers incorporates helper lipids with a phase transition temperature below room temperature (e.g., DMPC, DOPE), cationic lipids (e.g., DOTAP) for charge modulation, and PEGylated lipids (DSPE-PEG2000) for enhanced stability. We conducted an initial formulation screen with confocal microscopy to optimize fusion activity. The typical FLIP is unilamellar with surface charges that increase in proportion to the DOTAP ratio (**Fig.S1A, Fig.2A**). When incubated with a lung cancer cell line from a Kras^G12D^;Trp53^fl/fl^ (KP) mouse, FLIPs colocalized with the cell membrane (**Fig.2B**, purple in **Fig.2C**), and showed minimal overlap with endosomes and lysosomes (green in **Fig.2D)**. Conversely, non-fusogenic positively- (LP1) and negatively-charged (LP2) liposomes were primarily observed in locations overlapping with endosomes and lysosomes, as confirmed via Pearson Colocalization Coefficient (PCC) (**Fig.2D, E)**. These results indicate that FLIPs interact with cell membranes in a fusion-mediated manner, not through electrostatic adsorption or endocytosis. Moreover, we also found that membrane fusion propensity is a function of overall zeta potential and particle size (**Fig.S1**). Altogether, our screen suggested that FLIPs with 20% DOTAP, surface charge of ∼15mV, and ∼80 nm in diameter exhibited optimal fusion capability (**Fig.2A, Fig.S1B-D**), and were selected as the foundational formulation for protease-activatable C-FLIPs.

**Figure 2.**
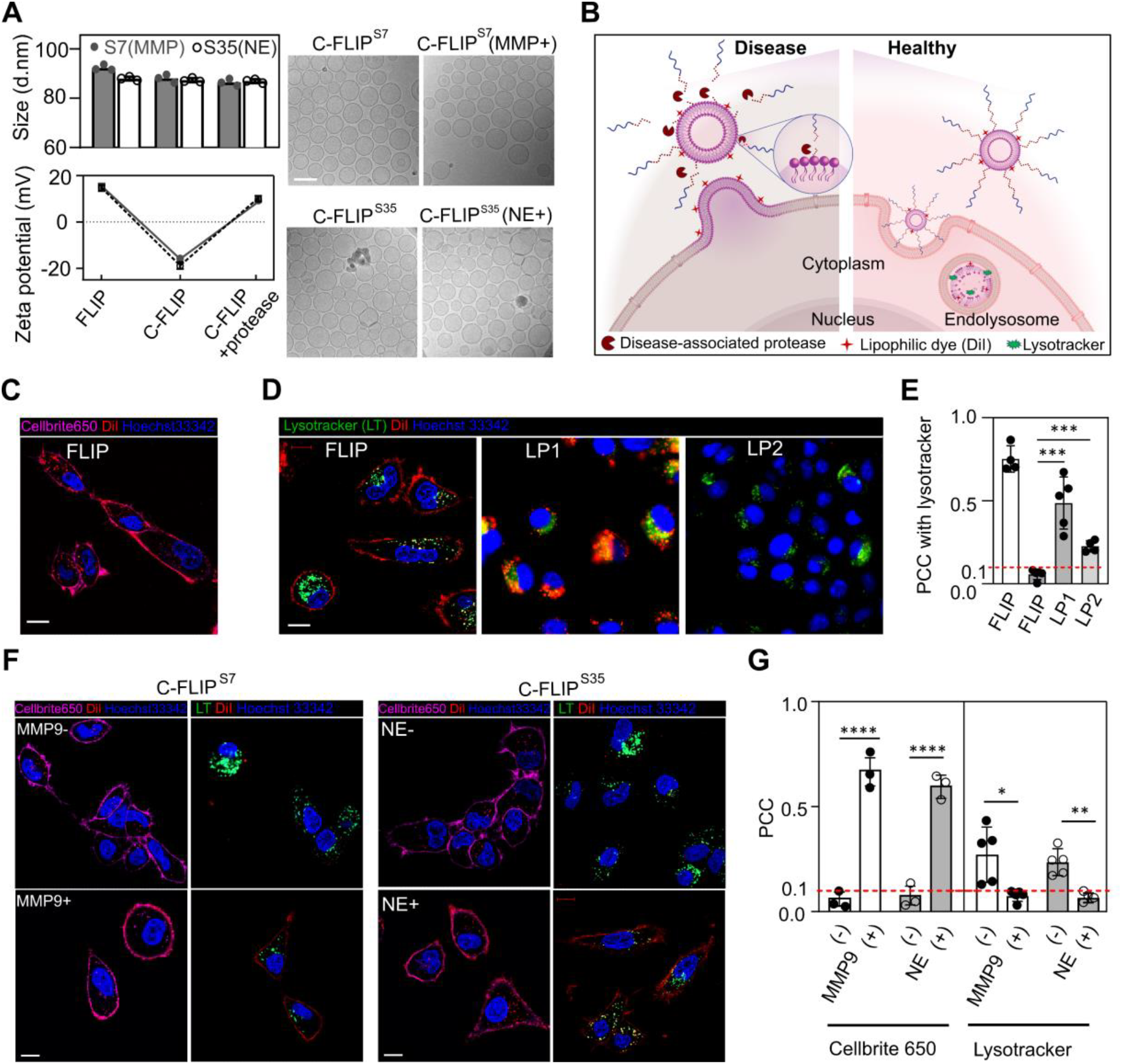
Protease activity orchestrates conditional fusion of liposomes with cells. **(A)** Size and zeta potential of FLIPs, C-FLIPs and protease pre-cleaved C-FLIPs. Peptide substrates S7 and S35 (refer to **Table S3** for the sequences) are preferentially cleaved by matrix metalloproteinase 9 (MMP9) and neutrophil elastase (NE), respectively. Poly(D-glutamic acid) (e8) was conjugated as a fusion mask. The scale bar of cryogenic transmission electron microscopy is 100 µm. **(B)** Scheme of intracellular trafficking of C-FLIPs in pathological and healthy microenvironment. Cognate proteases in the lesions cleave peptide substrates to liberate fusion masks, the C-FLIPs resume membrane fusion and enable lipophilic dye to label cell membrane, whereas masked C-FLIP is endocytosed and colocalized with lysotrackers in normal tissues. **(C)** Co-staining of cell membrane (Cellbrite 650) and FLIPs (Dil) in the mouse lung KP tumor (epithelial) cells with confocal microscopy. Scale bar = 5 µm. **(D)** Subcellular trafficking of FLIPs and non-fusogenic. LP1 and LP2 are positively and negatively charged, respectively. We labeled lipid layers of liposomes with lipophilic dye – DiI, nuclei with Hoechst 33342, endo/lysosomes with LysoTracker^TM^ DND-26 (LT), and cell membrane with Cellbrite 650. Scale bar = 5 µm. **(E)** Pearson correlation coefficient (PCC) of FLIP, LP1, and LP2 with cellular membrane (white bar) or endosomes/lysosomes (gray bars) in (C) and (D). \*\*\**p*<0.001 by one-way ANOVA with Dunnett’s multiple comparisons with FLIP. **(F)** Confocal microscopy of C-FLIPs with a fusion mask e8 conjugated with S7 peptides (C-FLIP^S7^) or S35 peptides (C-FLIP^S35^) before and after protease activation. The absence and addition of the matching proteases, MMP9 or NE, are indicated by ‘-’ and ‘+’, respectively. Scale bar = 5 µm. **(G)** PCC of C-FLIPs with cellular membranes (Cellbrite 650) and endosomes/lysosomes (lysotracker) in (F). \**p*<0.05, \*\**p*<0.01 and \*\*\*\**p*<0.0001 by Student’s t-test with respect to the protease-free group. All error bars represent standard deviations.

A core feature of C-FLIPs is the integration of a potent fusion mask that functions to prevent interactions with non-target cells. We nominated a variety of blocking agents that mask non-specific membrane interactions via surface charge shielding, such as negatively-charged peptides and oligonucleotides, or steric hindrance, such as bioinert polymers and zwitterionic molecules (**Table S1, Fig.S2A**). We conjugated these fusion mask candidates to maleimide-functionalized DOPE (DOPE-MAL) of the FLIPs through protease-cleavable peptide substrates via click chemistry. By incorporating optimal peptide substrates designed to be cleaved by disease-associated proteases, the liberation of fusion masks can be specifically triggered at target lesion sites to enable membrane fusion, while preventing fusion with non-target cells in systemic circulation and healthy tissues. We chose two peptide substrates, S7 and S35 (**Table S2, S3**) that can be cleaved by matrix metalloproteinase 9 (MMP9) and neutrophil elastase (NE), respectively, for use in *in vitro* analyses. These substrates were covalently coupled to a model negatively-charged fusion mask: poly(D-glutamic acid) octamer (e8), which we found to be the most effective at inhibiting fusogenic properties of FLIP in our initial screen (**Fig.S2B**,**C**). Additionally, e8 was incorporated into peptide substrates via a simple process during C-FLIP synthesis. While the size, polydispersity, and particle morphology remain unaltered upon conjugation, the surface charges of C-FLIPs drop to moderately negative, -15 to -20 mV (**Fig.2A**). We observed limited staining of the cell membrane by two C-FLIPs, and minimal internalization of the C-FLIPs without the addition of MMP9 or NE (top panel in **Fig.2F**), demonstrating that the conjugated e8 effectively suppressed membrane fusion of liposomes with KP cells. Upon *in vitro* incubation with MMP9 or NE, the positive charges of both C-FLIPs were nearly restored (**Fig.2A**), and accompanied by membrane fusion activity, as evidenced by re-staining of membrane and low PCC values with acidic subcellular compartments (**Fig.2F, G**). Note that the proportion of DOPE-MAL significantly influenced FLIPs surface charges as well as charge recovery after protease cleavage, even with a constant DOTAP ratio (**Fig.S1E**). We found that DOPE-MAL at 7% of all lipid components enabled the C-FLIPs to remain fully fusion-inactive until protease activation, but became fusion-efficient upon activation, outperforming other C-FLIPs with lower or higher DOPE-MAL ratios.

### Protease-mediated delivery of distinct classes of macromolecular therapeutics

We proceeded to assess whether the removal of blocking motifs (e.g., e8) from the C-FLIP by dysregulated extracellular proteases could enable fusion-mediated delivery of therapeutics into the cytosol. As a proof of concept, we studied the delivery of an impermeant small fluorescent molecule - calcein (0.6 kD), and a Cy5-labeled bovine serum albumin (BSA, 65 kD) encapsulated in NE- and MMP-cleavable C-FLIPs, respectively. When incubated with the KP cells for 1 h, we observed uniform distribution of calcein or BSA (**Fig.S3A-C**) across the cytoplasm if the C-FLIPs are pre-activated by the recombinant NE or MMP for 2 h that removes the e8 fusion mask. Without the addition of these recombinant proteases to *in vitro* cultured cells, the fusion mask e8 remains tethered to the liposomes, and consequently their payloads are largely delivered to endosomes and lysosomes, as measured by high Manders Colocalization Coefficient 1 (M1) that calculates the fraction of fluorescent marker that colocalize with endosomes and lysosomes (**Fig.S3D**). We expect that the differential expression of proteases present in biological contexts would dramatically increase *in vivo* efficiency of membrane fusion between C-FLIP and tumor cells, relative to fusion frequency occurring in healthy tissues and the systemic circulation.

We next evaluated functional readouts of therapeutic payloads delivered by C-FLIPs. Our initial model therapeutic was a peptide-based proteolysis-targeting chimera (pPROTAC) previously reported in the literature.^24^ Clinical translation of pPROTACs has been hampered by their impermeability into cells. Typically, cell-penetrating peptides (CPPs) are employed to translocate these pPROTACs into the cytoplasm. In our synthetic approach, we leveraged a protease trigger associated with underlying diseases to activate cytosolic delivery of the C-FLIPs. We utilized a MMP9-cleavable C-FLIP^S7^ to deliver a phosphorylated version of pPROTAC (termed ‘pP85’), reported to bind the P85 subunit of phosphoinositide 3-kinase (PI3K) and thereby promote proteasomal degradation (**Fig.3A**). Upon *in vitro* cleavage by MMP9, the pre-activated C-FLIP delivers phosphorylated pP85 and leads to ∼72% degradation of the P85 domain (**Fig.3B**). In contrast, CPP-tethered phosphorylated pP85 elicits only limited degradation of the target domain, likely due to the decrease in overall charge contributed by phosphorylation, which reduces its cell-penetrating capabilities.

**Figure 3.**
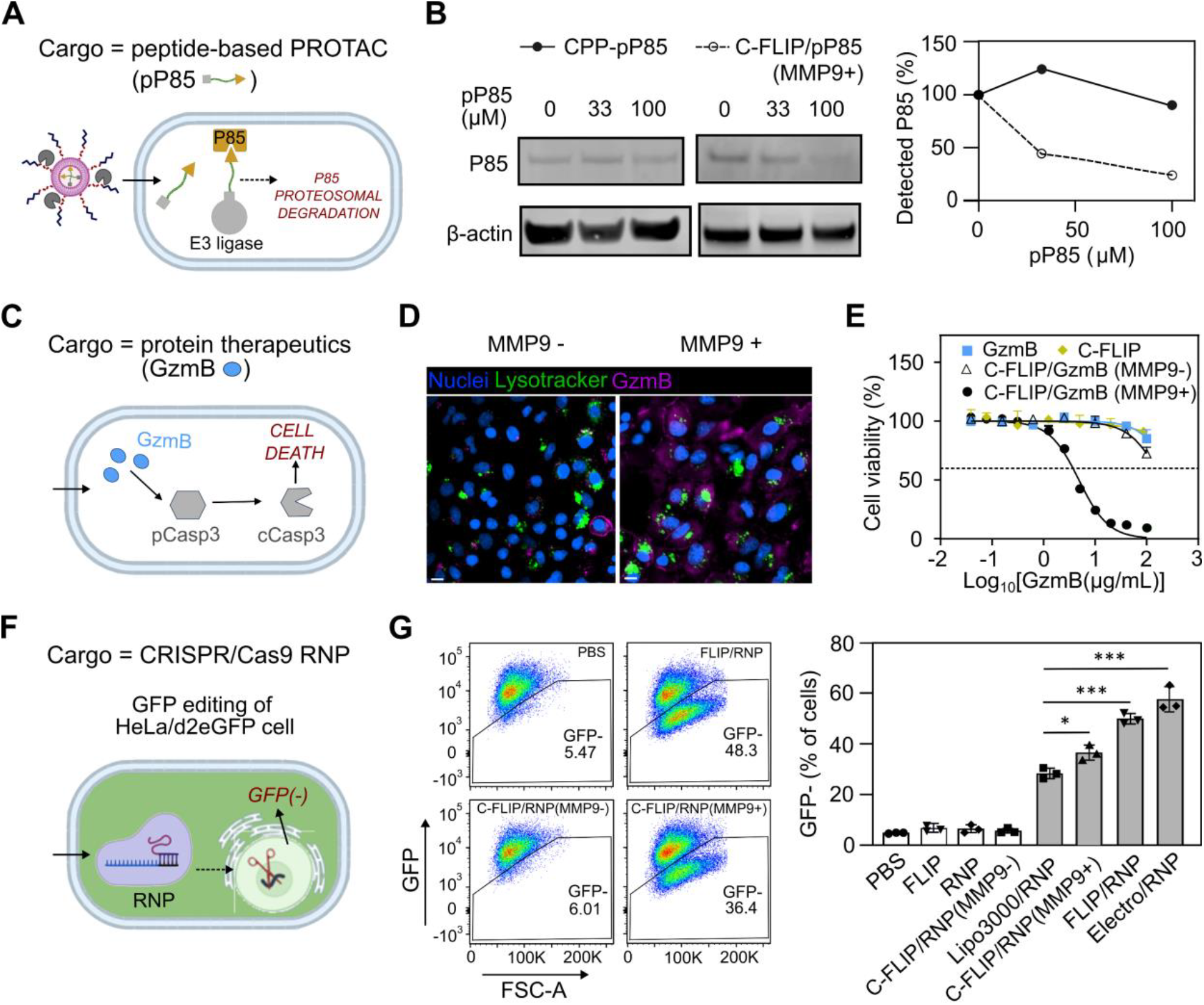
C-FLIPs enable direct cytosolic delivery of distinct macromolecular therapeutics in a protease-mediated manner. **(A)** Scheme of delivery of peptide-based proteolysis-targeting chimeras (pPROTAC) that couple target intracellular proteins, P85 domain of phosphoinositide 3-kinase (PI3K), to an E3 ligase (e.g, VHL) to facilitate ubiquitin (Ub) labeling of target protein for proteasomal degradation. pP85 = pPROTAC that targets the P85 domain. **(B)** Immunoblotting of the P85 domain of the KP cells treated with a phosphorylated pP85 that was delivered via pre-cleaved C-FLIP^S7^ [C-FLIP^S7^/pP85(MMP9+)] or tethered to a short cell-penetrating peptide (CPP) of poly(D-arginine) (CPP-pP85). **(C)** C-FLIP-mediated delivery of membrane-impermeable granzyme B (GzmB) to induce programmed cell death of target cells. **(D)** Cy5-labeled active GzmB (purple) is primarily delivered to the cytosol of the KP cells with C-FLIP^S7^ in the presence of recombinant MMP9(+), while the un-cleaved C-FLIP^S7^ localizes the GzmB in endosomes and lysosomes (green). Scale bar = 5 µm, **(E)** Cell viability (24 h) of the KP cells treated with the pre-cleaved C-FLIP^S7^/GzmB or control groups including C-FLIP ^S7^/GzmB in absence of MMP9(-). **(F)** Delivery of ribonucleoprotein particles (RNPs) of a recombinant CRISPR-Cas9 enzyme and guide RNA targeting destabilized GFP with pre-cleaved C-FLIP^S7^ (MMP9+) in HeLa cells expressing destabilized GFP (HeLa/d2eGFP). **(G)** Efficient GFP knockout (GFP-) by FLIP or pre-cleaved C-FLIP delivery of RNPs (Cas9 50 µM; n=3 per group). GFP knockout efficiency was quantified with flow cytometry and benchmarked against commercial Lipofectamine 3000 (Lipo3000/RNP) transfection reagents and electroporated RNPs (Electro/RNP). \**p*<0.05, \*\*\**p*<0.001 and \*\*\*\**p*<0.0001 by one-way ANOVA with Dunnett’s multiple comparisons with respect to Lipo3000/RNP. All error bars represent standard deviations.

We then utilized the same C-FLIP^S7^ to encapsulate a much larger (30kD) cationic protein – GzmB (**Fig.S4A, B**), a serine protease secreted by activated T cells and natural killer cells that induces programmed cell death via activation of caspases and other intracellular targets (**Fig.3C**). To test for loading efficiency, we first compared sonication and repeated freeze-thaw for encapsulating GzmB in C-FLIP and found that GzmB-mediated catalysis remained intact after 10 freeze-thaw cycles but dropped nearly 30% with as little as 1 minute of mild sonication (**Fig.S4C-F**). Therefore, we opted for freeze-thaw cycles to encapsulate protein-based payloads. Fluorescent microscopy was used to track the subcellular localization of Cy5-labeled GzmB. As predicted, pre-activation of C-FLIP^S7^/GzmB by MMP9 resulted in cytosolic distribution of GzmB (**Fig.3D**). The KP lung cancer cells were killed effectively by GzmB delivered via pre-activated C-FLIP, but not by empty C-FLIP^S7^ vehicle, naked GzmB, or non-activatable N-FLIP^S7^/GzmB (**Fig.3E**). The caspase 3 activity was detected intracellularly (**Fig.S4G**). Similar *in vitro* killing efficacies were observed in multiple mouse cancer cell lines (**Fig.S4H**).

Finally, we explored the utility of C-FLIP to enable the conditional cytosolic delivery of genome editors in RNP complexes. Cas9 enzyme and single-guide RNA (sgRNA) targeting a destabilized green fluorescent protein (deGFP) were formed into RNP complexes of 15-20 nm in diameter with a slightly negative surface charge (**Fig.S5A, B**). Upon encapsulation, the Cas9-gRNA RNP delivered via either unmodified FLIP or pre-activated C-FLIP^S7^ led to a substantial reduction of GFP expression in a deGFP+ HeLa cell line (**Fig.3F**). In comparison to cells treated with a commercial transfection reagent, Lipofectamine 3000, which achieved a ∼30% reduction, both FLIP (∼50%) and pre-activated C-FLIP (∼40%) formulations significantly outperformed the current standard (**Fig.3F, Fig.S5C**).

### Dysregulated proteases in the tumor microenvironment enhance on-target delivery and tolerability of fusogenic liposomes

To evaluate disease site-specificity for targeted delivery of C-FLIP in cancer applications, we established a lung metastasis tumor model by inoculating mice with the KP cancer cell line via tail vein,^25^ and subsequently assessed disease-associated protease activity. We screened a library of 39 Förster resonance energy transfer (FRET)-paired peptides against lung homogenates collected from the tumor-bearing mice 14 days after inoculation. We found that 15 peptide substrates were differentially cleaved compared to incubation with homogenates from healthy mice. Notably, S7 showed a ∼4-fold increase in cleavage activity in tumor samples (**Fig.S6**), suggesting its potential to selectively activate C-FLIP in the tumor microenvironment (TME). In comparison, S24 showed a moderate increase (∼1.8-fold, *p*<0.05 by Student’s t-test), whereas the cleavage profile of S35 was similar in both healthy and tumor-bearing lungs. Our previous work showed S7, S24, and S35 were preferentially cleaved by MMP9, urokinase-type plasminogen activator (uPA), and NE, respectively.^26,27^

The conjugation of e8 onto the FLIP via S7 or S35 prolonged circulation half-life by 20-fold, relative to unmodified FLIP in lung tumor-bearing mice (**Fig.4A**). We hypothesized the extended circulation may enhance the likelihood for disease-specific proteases to perform catalytic reactions on the C-FLIPs. While cationic DOTAP is reported to increase the transport of lipid nanocarriers towards the lungs,^28^ we observed limited lung accumulation of the FLIP (<2% of injected dose) in the mice with KP lung tumors (**Fig.4B and 4D**), indicating the altered therapeutic profile with peptide conjugation. The choice of protease substrates in C-FLIPs was crucial, serving not only for conditional activation but significantly influencing fusogenic nanoparticle biodistribution. We compared the biodistribution of 3 C-FLIP formulations, using either S7, S24, or S35 as the peptide substrates for the conjugation of conditional mask to FLIP. Approximately 8% of injected MMP9-cleavable C-FLIP^S7^ accumulated in the tumor-bearing lungs, which was ∼2 fold higher than C-FLIP^S24^. We further noted that C-FLIP^S35^ did not show significant accumulation in lung tumor nodules. C-FLIP^S7^ demonstrated more than 3-fold higher accumulation than C-FLIP^S35^. Noticeably, less than 2% of the MMP9-cleavable C-FLIP^S7^ accumulates in the lungs of normal mice. These findings highlight the importance of protease activity-driven tumor tropism and peptide substrate selection. To further enhance tumor homing and penetration, we sought to optimize C-FLIP by incorporating a prominent tumor-homing peptide, iRGD,^29,30^ designated as TC-FLIP^S7^. This modification results in nearly 15% of the liposomes trafficking to tumor-bearing lungs, as well as improved pharmacokinetics (**Fig.4C and 4D**).

**Figure 4.**
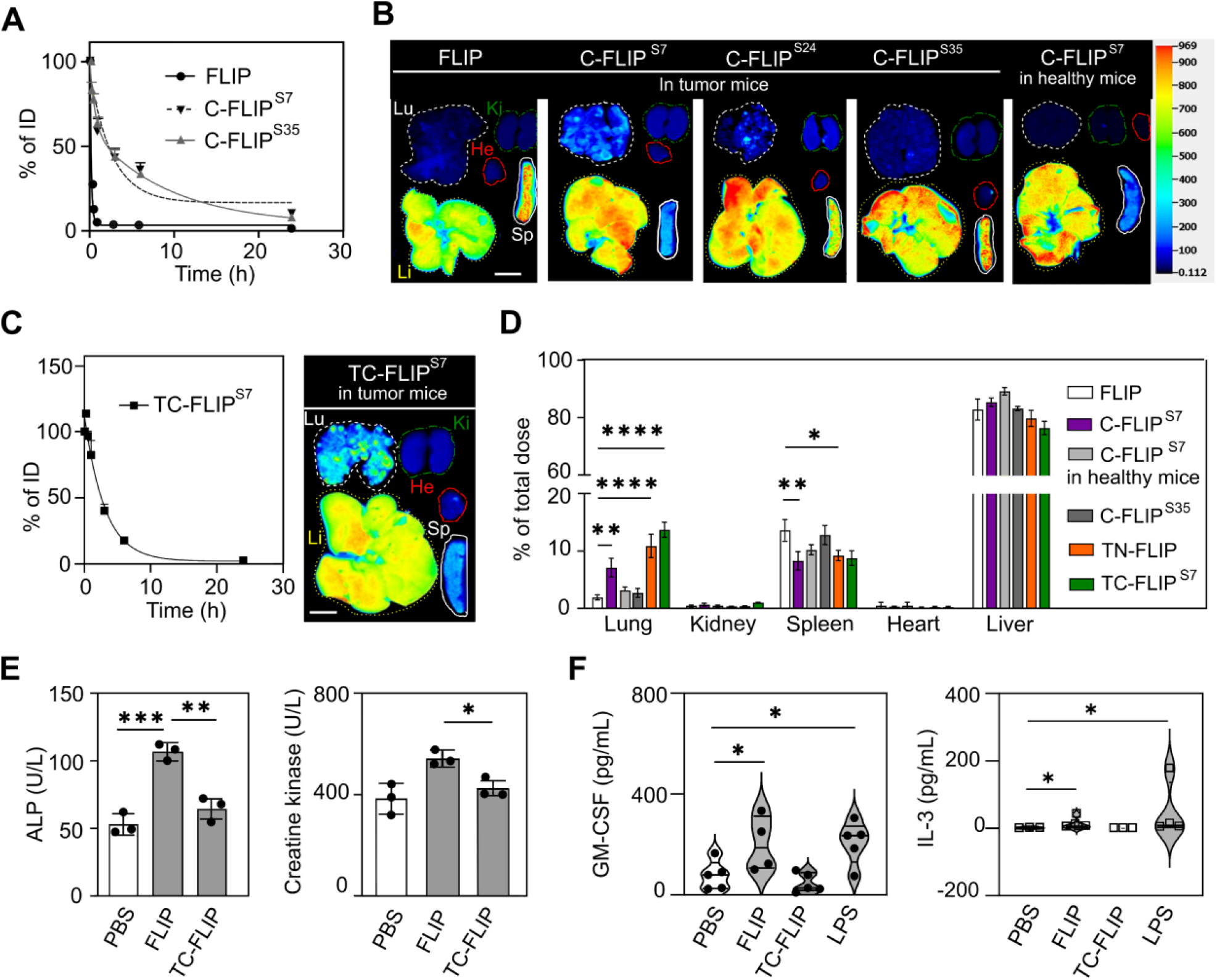
Tumor-specific protease activity and targeting ligands combine to improve tumor tropism of C-FLIPs. **(A)** Plasma clearance and half-life time (t_1/2_) of intravenously-injected (i.v.) FLIP and C-FLIPs. ID = injected dose. t_1/2_ of FLIP, C-FLIP^s7^ and C-FLIP^s35^ are 0.11, 1.84 and 2.27 h. **(B)** Representative images of 24 h biodistribution of i.v. FLIP and C-FLIPs in the mice with KP lung tumors. Tumor-specific cleavage of S7 and S24 significantly increases the C-FLIP accumulation in the lung tumor nodules, while tumor non-specific cleavage of S35 minimally alters liposomal accumulation in lung tumor nodules. Lu=lung, Li=liver, He=heart, Ki=kidney and Sp=spleen. Scale bar = 1 cm. **(C)** Tumor homing peptide, iRGD, can further augment tumor targeting of C-FLIP^S7^(TC-FLIP^S7^), with half-life time (t_1/2_=2.12 h) remaining similar to the C-FLIP counterparts. **(D**) Quantifying 24 h biodistribution of i.v. FLIP and TC-FLIPs in the KP lung tumor-bearing and healthy mice. TN-FLIP = targeted, non-cleavable FLIP with fusion mask e8 conjugated directly to the FLIP. \**p*<0.05, \*\**p*<0.01 and \*\*\*\**p*<0.0001 by one-way ANOVA with Dunnett’s multiple comparisons in comparison to the FLIP. All error bars represent standard deviations. Fluorescence intensity ladder and scale bar are same as (B). **(E)** Toxicity of FLIP and TC-FLIP assessed in healthy wild-type C57BL/6 mice by general clinical chemistry. A panel of 23 serum analytes were examined (**Fig.S7**) and only alkaline phosphatase (ALP) and creatine kinase are presented here as they show significant difference between the FLIP and C-FLIP^S7^. **(F)** Systemic immune responses elicited by the FLIP and TC-FLIP. Thirty-two serum cytokines were analyzed in healthy C57BL/6 mice (**Fig.S7**). Granulocyte macrophage-colony stimulating factor (GM-CSF) and interleukin 3 (IL-3) show significant increase after the FLIP exposure.

In addition to enhancing the biodistribution profile, e8 improved the safety relative to unmodified FLIPs. Multidose administration of TC-FLIP did not cause weight loss in normal wild-type mice (**Fig.S7A, B**). Lower serum levels of both alkaline phosphatase (evaluating liver function) and creatine kinase (heart function) were also observed (**Fig.4E, Fig.S7D)**. Through H&E staining, we did not observe any acute toxicity with multidose of intravenously injected TC-FLIP **(Fig.S7E)**. The TC-FLIP formulation also did not increase the levels of circulating cytokines (e.g., IL-3, GM-CSF in **Fig.4F and Fig.S7C**) associated with immunogenicity. Given the superior tumor targeting, improved safety profile and high cleavage efficiency by tumors, we nominated TC-FLIP as our optimal delivery platform for subsequent *in vivo* validation of antitumor efficacy.

### GzmB delivered by C-FLIP induces immunogenic cell death via gasdermin E-mediated pyroptosis

Despite its promise, cancer immunotherapy has been confronted with limitations attributed to an immunosuppressive (‘cold’) microenvironment. We hypothesized that engineering a tumor-targeted TC-FLIP that enables cytosolic delivery of GzmB to mimic functional cytotoxic T cells might sensitize cold tumors to immune therapy. We encapsulated GzmB in TC-FLIP and measured their impact on pro-inflammatory pathways *in vitro* and *in vivo*. Using an initial *in vitro* assay, we treated KP lung cancer cells and examined the expression of immunogenic cell death (ICD) markers: calreticulin (CRT), high mobility group box 1 (HMGB1), adenosine triphosphate (ATP), and interleukin-1β (IL-1β) on the cells or in the culture media (**Fig.5A**). Activated TC-FLIP/GzmB induced the highest levels of externalized CRT, release of HMGB1, and ATP after a 24 h treatment, compared to naked GzmB, empty FLIP vehicle, or non-activatable GzmB-loaded TN-FLIP/GzmB (**Fig.5B-D**). However, IL-1β was not activated in any of treated groups (**Fig.5E**). Interestingly, TC-FLIP/GzmB results in a bimodal distribution of CRT surface exposure (purple line in **Fig.5B**), indicating a mixture of apoptotic and pyroptotic cells. Additionally, TC-FLIP/GzmB treatment gave rise to N-terminal proteolytic fragments of gasdermin E (GSDME) (**Fig.5F**), indicating non-canonical pyroptosis via a GzmB-caspase 3-GSDME axis, without gasdermin D (GSDMD) activation (**Fig.5F**) or active IL-1β (**Fig.5E**). Such an inflammatory form of programmed cell death holds the potential to sensitize immunologically cold tumors to immune checkpoint inhibitors (ICIs).

**Figure 5.**
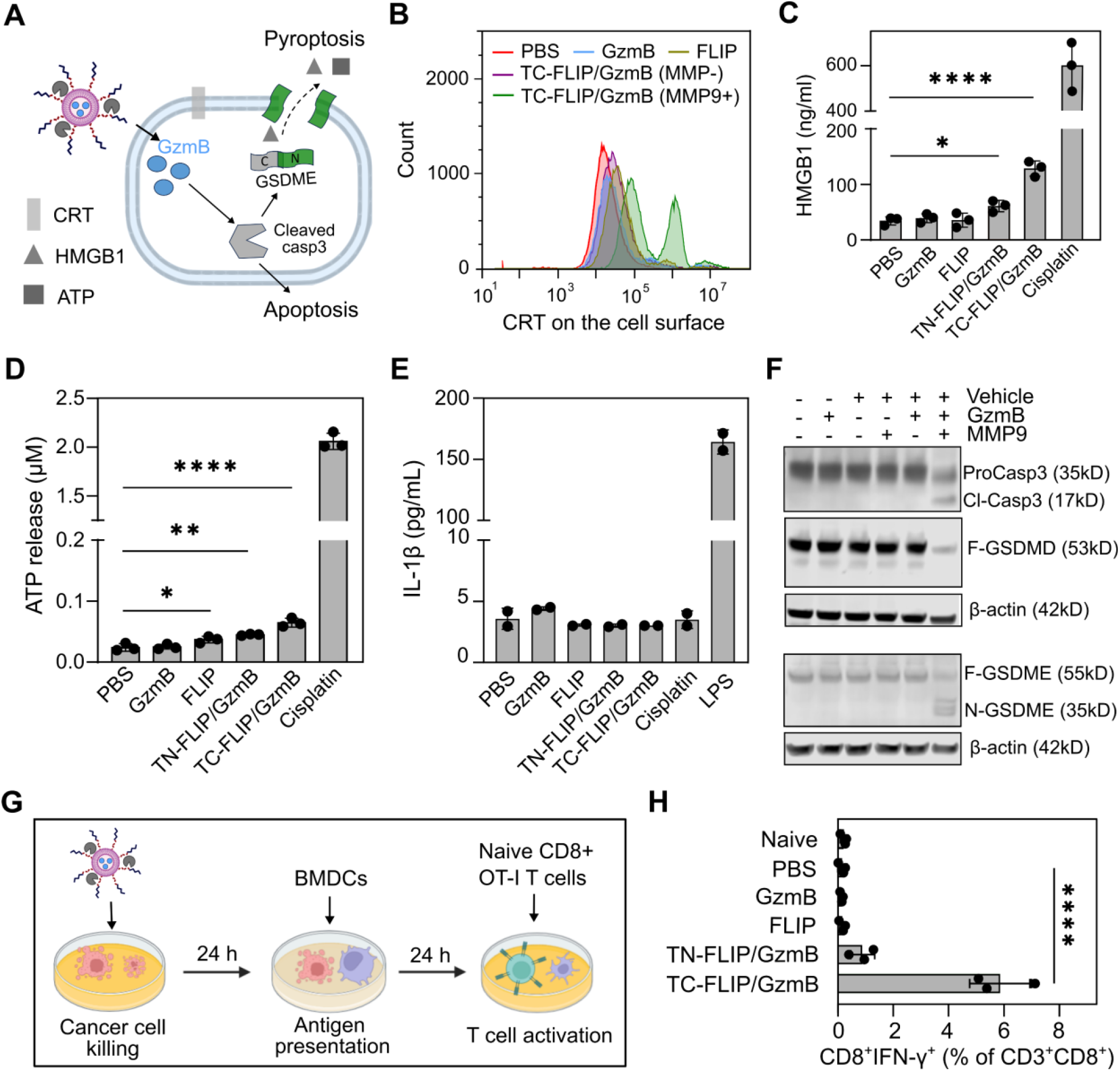
C-FLIP/GzmB promotes immunogenic cell death via gasdermin E-mediated pyroptosis. **(A)** Granzyme B (GzmB) not only induces immunologically-inert apoptosis, but also activates both caspase 3 (casp3)-dependent and -independent pyroptosis in target cells that highly express gasdermin E (GSDME), such as KP cells. The formation of transmembrane pores by gasdermin promotes the leakage of typical immunogenic cell death (ICD) markers: calreticulin (CRT), high mobility group box 1 protein (HMGB1) and adenosine triphosphate (ATP). **(B)** flow cytometry detects CRT on the surface of KP cells. **(C)** HMGB1, **(D)** ATP, but not **(E)** interleukin-1 beta (IL-1β) were detected in the culture media of KP cells treated with the pre-cleaved TC-FLIP/GzmB (MMP9+) for 24 h. Cisplatin was used as a positive control to induce the release of HMGB1 and ATP, while lipopolysaccharide (LPS) was used to activate the pro-form of IL-1β. **(F)** Immunoblotting detects one 19kD cleaved caspase 3 (Cl-Casp3) domain, pore-forming N terminus of GSDME (N-GSDME), but not pore-forming N terminal domain of GSDMD in the KP cancer cells treated with the pre-cleaved TC-FLIP/GzmB. None of these targets was detected in other treatment groups. β-actin was used as a protein loading control. ‘-’ and ‘+’ denote whether the component is excluded or included, respectively. ProCasp3 = pro-caspase 3. **(G)** *In vitro* antigen cross-presentation among TC-FLIP(GzmB)-treated B16-OVA cells, bone marrow-derived dendritic cells (BMDCs) and naïve OT-1 CD8+ T cells. TC-FLIP/GzmB induces surface presentation of antigen OVA from engineered B16 cells, which matures BMDCs after 24 h incubation. Naïve OVA-specific T cells are then activated by mature dendritic cells to produce antitumor cytokines. **(H)** Quantification of interferon-γ (IFN-γ)-producing CD3+CD8+ T cells to which dendritic cells were cross-presented OVA antigen. For all the above panels, error bars indicate standard deviations. \**p*<0.05, \*\**p*<0.01, \*\*\**p*<0.001 and \*\*\*\**p*<0.0001 by one-way ANOVA with Dunnett’s multiple comparisons with respect to the PBS group.

We then utilized another GSDME-positive cell line, B16 mouse melanoma engineered to express ovalbumin (B16-OVA), to study the interaction between pyroptotic cancer cells and T cells. We treated the B16-OVA cells with pre-cleaved TC-FLIP/GzmB, and examined its capability to induce dendritic cell (DC)-mediated antigen cross-presentation to naïve, OVA-specific CD8+ T cells. After 24 h treatment, the B16-OVA cells (including both injured and dead cells) were co-cultured with bone marrow–derived DCs (BMDCs) for an additional 24 h, after which purified, naïve CD8+ OVA-TCR transgenic (OT-1) splenic T cells were added to the coculture. We then measured activation of the OVA-specific CD8+ T cells, and found that only TC-FLIP/GzmB-treated B16-OVA effectively elicited significant levels of interferon-γ (IFN-γ) production in CD8+ T cells (**Fig.5G**). Furthermore, the culture media isolated from the B16-OVA/BMDC coculture was examined for DC activation. TC-FLIP/GzmB-treated cells upregulated typical co-stimulatory molecules with significance (**Fig.S8B-F**): CD86 (∼3.8 fold, \*\*\*\**p*<0.0001), CD80 (∼3.5 fold, \*\*\*\**p*<0.0001), CD40 (∼2 fold, \*\*\**p*<0.001), and pMHC-II (∼1.8 fold, \**p*=0.03) in naïve BMDCs, indicating DC maturation was induced by released tumor-associated antigens and ICD biomarkers, and thus would likely enhance T cell activation and priming in the TME and draining lymph nodes.

### TC-FLIP/GzmB sensitizes tumors to checkpoint blockade

Finally, we examined the efficacy of TC-FLIP/GzmB as an anti-tumor therapy *in vivo*. To establish tumors, the mice were inoculated with luciferase-expressing KP cells via tail vein injection, and treated with different regimens from Day 7 to 16 (**Fig.6A**). Treatment with only TC-FLIP/GzmB substantially suppressed lung tumor growth, as measured *ex vivo* by luminescence **Fig.6B, Fig.S9A**, histologically in sections (**Fig.S9C**), and via expression of proliferative markers (**Fig.S10**). Notably, the combination treatment resulted in a survival advantage (**Fig.S9B**), outperforming both non-specific FLIP/GzmB and the non-activatable masked formulation TN-FLIP/GzmB. Furthermore, TC-FLIP/GzmB demonstrated comparable antitumor potency to αPD-1 treatment, an established ICI therapy (**Fig.6B, Fig.S9, Fig.S10**). However, neither C-FLIP/GzmB nor αPD-1 was able to shrink KP tumors in any treated individuals. Strikingly, the combination of TC-FLIP/GzmB with αPD-1 led to tumor regression and even complete response (i.e., undetectable nodules with microCT) in some mice, suggesting that TC-FLIP/GzmB would substantially potentiate the ICI efficacy in the immunologically cold KP lung tumors^25^ that express considerable levels of GSDME (**Fig.S11A**). To test this hypothesis, we repeated these experiments in a model of lung metastasis based on the mouse colon cancer line, CT26, which expresses low levels of GSDME **(Fig.S11B**). The impact of the combined TC-FLIP/GzmB with αPD-1 treatment was less robust in the CT26-derived lung metastasis (**Fig.S12**), which is consistent with our hypothesis and suggests a path for tumor stratification in patients given ICI therapy. We then sought to characterize *in vivo* activation of caspase 3 in the KP tumor. Immunofluorescent staining revealed the elevated pro-caspase 3 cleavage in KP lung tumor nodules of TC-FLIP/GzmB- or αPD-1-treated groups, with even higher levels for the group receiving the combination treatment (**Fig.6C, Fig.S13A**). Noticeably, cleaved casp3 highly colocalized within cancer cells (up to 80% of CK8+ cells) (**Fig.S13B**).

**Figure 6.**
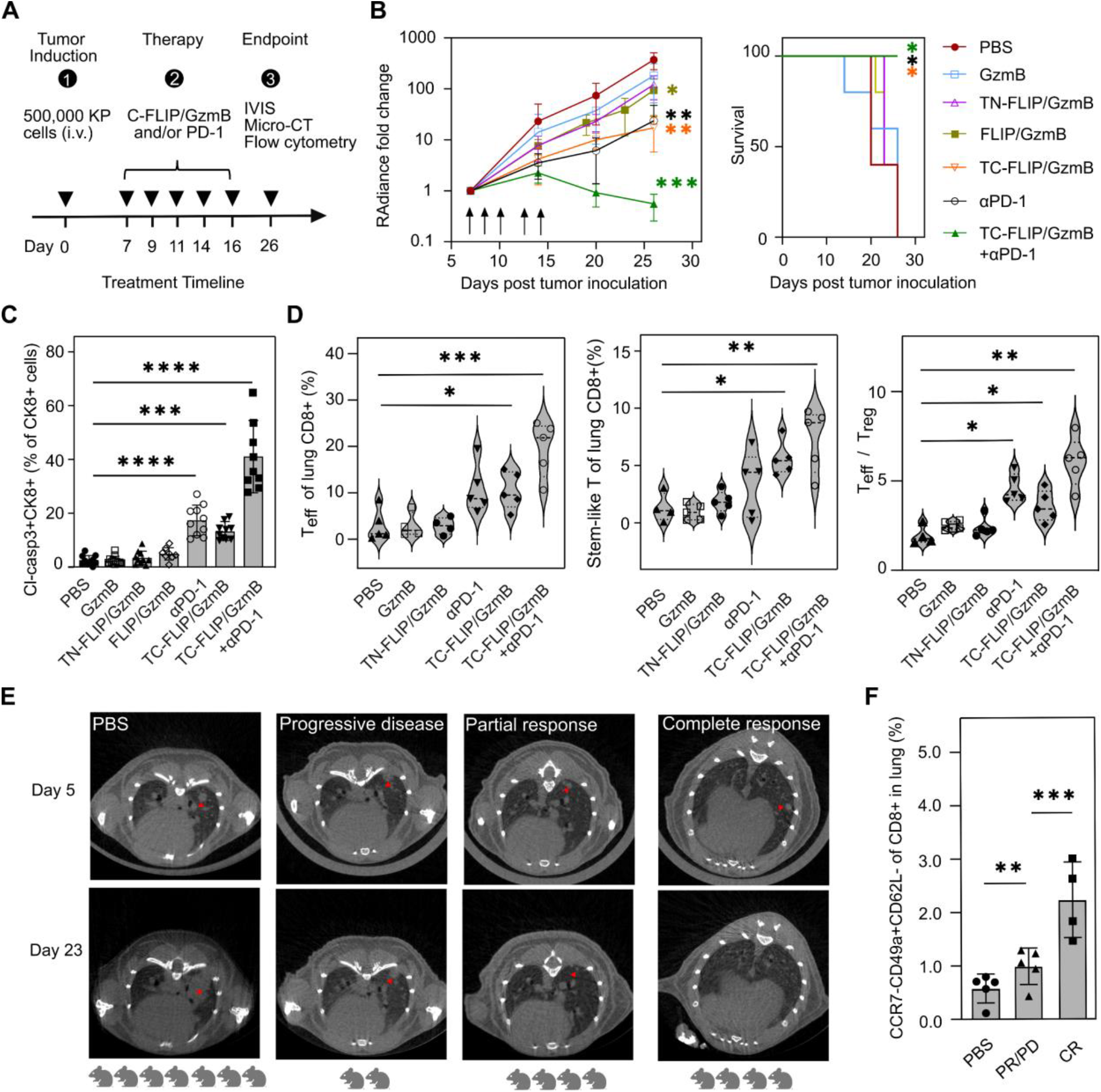
TC-FLIP/GzmB synergizes with PD-1 checkpoint inhibitor and tunes the tumor immune microenvironment. **(A)** Timeline for therapeutic efficacy study in a KP lung metastatic tumor model. Luciferase-transfected KP lung cancer cells were injected intravenously (i.v.) via tail vein. Treatments were administered on Day 7, 9, 11, 14 and 16 (red triangles) for 5 doses and tumor burden was regularly monitored via IVIS. In the combination group, C-FLIP/GzmB and PD-1 checkpoint inhibitor were administered on the same day (red arrows). **(B)** Longitudinal monitoring of tumor burden and survival curve of the mice treated with different regimens. **(C)** Fraction of dying cancer cells (Cl-Casp3+CK8+). Cl-Casp3 = cleaved caspase 3, CK8=cytokeratin 8. Flow cytometric quantification of the fraction of **(D)** effector T (T_eff_) cells (CD44+GzmB+CD8+) among lung CD8+ population, fraction of proliferative stem-like (progenitor) CD8+ T cells in the tumor-bearing lungs of different treated groups, and the ratio of T_eff_ cells to regulatory T (T_reg_) cells (CD4+CD25+Foxp3+). **(E)** Representative microCT images of lung tumors (red arrows) on Day 5 post tumor inoculation and Day 23 (one week post last treatment). The tumor-bearing mice were treated with PBS or TC-FLIP/GzmB+αPD-1 starting on Day 5 for 6 doses. The responses to TC-FLIP/GzmB+αPD-1 were categorized into progressive disease (PD), partial response (PR), and complete response (CR). The number of cartoon mice represents the size of PBS, PD, PR and CR groups. The mice from PBS group were scanned on Day 14 due to faster tumor growth. **(F)** Phenotypic analysis of resident memory T cells in the lungs of mice that responded to (CR) or resisted (PD+PR) TC-FLIP/GzmB+αPD-1 treatment from (E). PBS group was harvested for lungs on Day 16. One non-responder from PD group died before the terminal point. \*\**p*<0.01 and \*\*\**p*<0.001 by one-way ANOVA with Dunnett’s multiple comparisons with respect to PBS group.

To understand whether TC-FLIP/GzmB could potentially tune the tumor immune microenvironment (TIME) in the KP lung tumor model *in vivo*, we performed phenotypic profiling of T cells in the lungs. TC-FLIP/GzmB treatment increased the fraction of effector T (T_eff_) cells (CD44+GzmB+) among the lung CD8+ population by nearly 2.5 folds, comparable to the level observed in the αPD1-treated group, while the group administered the non-cleavable version, TN-FLIP/GzmB, maintained a similar population of effector T cells as seen in the PBS group. As expected, the combination of TC-FLIP/GzmB and αPD-1 resulted in the highest increase of effector T cells (**Fig.6D, Fig.S14A, B**). The fractions of stem-like, proliferative CD8+ T cells (CD44+TCF-1+Ki67+) also significantly increased in the TC-FLIP/GzmB (∼3.5 fold), αPD-1 (∼2.5 fold), and the combination groups (∼5.5 fold) (**Fig.6D, Fig.S14C**). These stem-like precursors harbor high self-renewal and proliferative capability, and are critical to confer sustained antitumor responses.^31^ In addition, TC-FLIP/GzmB as monotherapy and the combination of TC-FLIP/GzmB with αPD-1 both elevated the ratio of T_eff_ to regulatory T cells (CD4+CD25+Foxp3+), suggesting that immunosuppression had been reduced in the TIME (**Fig.6D**). We also evaluated the exhaustion status of CD8+ T cells in the TIME and found that both αPD-1 monotherapy and TC-FLIP/GzmB+αPD-1 combination significantly delayed the exhaustion of CD8+ T cells (PD-1+TIM-3+), whereas treatment with TC-FLIP/GzmB alone slightly altered the exhaustion of effector T cells (**Fig.S14D, E**). Collectively, these variations suggest that targeted, conditional FLIP-mediated delivery of GzmB could potentially inflame the TIME and significantly boost the efficacy of PD-1 blockade in immune cold tumors. Furthermore, we observed that full responder (CR) mice had higher levels of lung-resident memory CD8+ T cells after combinatorial treatment (**Fig.6E**) than did mice with partial response or little/no response (**Fig.6F**), suggesting enhanced long-term immune protection.

## Discussion

Recent advances in chemical biology, genome editing, and protein engineering have significantly expanded the application of biomacromolecules that are intended to function in the cytoplasm. However, their effectiveness is often hampered by their rapid excretion, systemic degradation and entrapment in endosomes. Encapsulation of therapeutic molecules in non-viral fusogenic vehicle that bypasses endocytosis has demonstrated potential in addressing these challenges. Nonetheless, many of these fusogenic delivery systems still suffer from a lack of spatiotemporal specificity in the pathological context and pose risks of off-target toxicity. Here, we engineered C-FLIPs that exploit tumor-specific, dysregulated extracellular proteases to redirect nanoparticles to extra-hepatic disease sites and subsequently unveil the fusion-mediated cytoplasmic delivery of biologics (**Fig.1**). Using this platform, masked C-FLIPs remain therapeutically inactive in healthy tissues to minimize off-target effects. The defining aspect of C-FLIPs is the incorporation of fusion masks via protease-labile peptide linkers that effectively block their membrane fusion (**Fig.2**). The fusion masks could function via mechanisms such as surface charge shielding, steric hindrance, and/or their combination. Although the results indicate that the negatively-charged peptide (e8) is the most effective in fusion suppression and offers a one-pot synthetic advantage, other fusion masks explored also show promise and may benefit from further optimization and evaluation of collateral functions (e.g., synergistic therapies, diagnostic readouts, disease targeting). In addition to restricting fusion, the negatively-charged fusion mask e8 not only prolongs C-FLIP circulation time by neutralizing surface charges (**Fig.4A**), but also enhances the safety profile of this nanoparticle delivery system (**Fig.4E, F**). Therefore, leveraging conditional fusion has the potential to enhance selective cytosolic delivery via fusogenic lipid-based vehicles and virus-like particles (VLPs) under a rapid development.^15,20–22,32^

Protease-labile peptide substrates that serve as triggers of C-FLIP fusogenicity, offer substantial advantages over other endogenous (e.g., pH, hypoxia) and external stimuli (e.g., ultrasound, laser irradiation) due to their abundance, flexibility, accessibility, and selectivity.^33–36^ By leveraging disease-specific protease activity—such as type IV collagenases in solid tumors—C-FLIPs achieve precise, conditional delivery, selectively fusing with cell membranes in distinct disease environments. Proteases play functional roles in driving disease progression, and display distinct expression patterns and catalytic activity signatures across hallmarks (e.g., survival, angiogenesis, metastasis) and stages of diseases, such as cancer.^36–39^ These distinct proteases are spatially located in the TME and/or associated with cells as soluble, matrix-bound, or membrane-bound forms. By profiling these disease-specific proteases and employing advanced substrate design (e.g., OR logic-gated sequences, machine learning, high throughput screening), we can fine-tune C-FLIPs for precise activation, significantly reducing adverse off-target effects. Furthermore, the capacity to design bespoke C-FLIP sequences may enable options for personalized therapy,^40,41^ as well as to develop an atlas of highly modular C-FLIPs that establish a “plug-and-play’ approach to tackle various disease conditions. Altogether, protease activities offer a modular and precise toolkit that can unveil conditional fusion constructs in a spatiotemporal manner, compared to existing fusogenic lipid-based nanoparticles, virus-like particles, and their activatable counterparts that rely on individual microenvironmental triggers (e.g., reactive oxygen species).^42^

In this study, we have successfully demonstrated C-FLIP versatility to encapsulate not only peptides and enzyme therapeutics, but also RNPs for cytosolic delivery (**Fig.3**), expanding their utility in delivering genome editing tools. Delivering RNPs with C-FLIPs could offer another advantage in reducing off-target effects, compared to mRNA-based methods, owing to the shorter duration of genome exposure to the RNPs. By leveraging abnormal protease expression and activity in candidate genetic diseases, we would anticipate that the C-FLIP/RNP platform could be used for therapeutic gene correction while minimizing off-target effects, as well to reduce incorrect gene and RNA editing of non-target cells.

Our demonstration of the therapeutic efficacy of a GzmB-encapsulating C-FLIP, TC-FLIP^S7^/GzmB, in a cancer model was used as a proof of concept regarding the potential impact of this approach. We observed strong antitumoral potency in a Kras-mutated, p53-inactivated mouse lung tumor model. The key highlight is that the C-FLIP/GzmB could access the tumor core and initiate GSDME-mediated pyroptosis, rather than apoptosis, in tumor cells. This change inflames the otherwise immunosuppressive TME, and enhances the efficacy of immune checkpoint PD-1 inhibitors. The synergy between C-FLIP/GzmB and ICIs is prominent in lung tumors derived from a KP cell line that highly expresses GSDME, but to a lesser degree in CT26-derived lung metastasis that exhibits very low GSDME expression (**Fig.6B, Fig.S11**). In future, a GSDME knockout mouse model could further elucidate the pivotal role of GSDME in the antitumor response of C-FLIP/GzmB. GSDME often undergoes cleavage site mutation and subsequently is less activated in many cancer cells,^43^ thus we anticipate that co-delivery of exogenous GSDME and GzmB with TC-FLIP may serve as a universal approach to inflame cold tumors, regardless of T cell infiltration and expression of endogenous GSDME.

Looking ahead, the application of C-FLIPs across diseases beyond the realm of cancer presents an opportunity to substantially increase their impact. Diseases characterized by unique protease profiles, including various infections, inflammatory, and degenerative conditions, could greatly benefit from the precise delivery capabilities of C-FLIPs. The multifaceted functionality of C-FLIP highlights its potential as a transformative tool in precision medicine, offering new avenues for therapeutic intervention. The modularity of C-FLIP allows for the customization of its particle properties, enabling targeted, efficient, and safe treatment delivery across a wide spectrum of medical conditions. These are key factors in enhancing patient outcomes and quality of life. Broadly, the non-viral, cell-free, highly modular C-FLIPs represent a promising platform that enables targeted cytosolic delivery of membrane impermeable therapeutics, and could provide new tools of precision medicine in immunotherapy and gene editing.

## Acknowledgments

We are grateful to the Koch Institute Swanson Biotechnology Center, specifically the Histology core, the Biopolymer and Proteomics core, and Nanotechnology Materials core. Funding: This work was supported by the Koch Institute’s Marble Center for Cancer Nanomedicine, the Koch Institute Frontier Research Program via the Kathy and Curt Marble Cancer Research Fund, and the Virginia and D.K. Ludwig Fund for Cancer Research. Additional funding provided by a Koch Institute Support (core) Grant P30-CA14051 from the National Cancer Institute and a Core Center Grant P30-ES002109 from the National Institute of Environmental Health Sciences. S.N.B. is an HHMI investigator. E.K.W.T received a postdoctoral fellowship from the Ludwig Center at MIT and support via the Convergence Scholars Program from the Marble Center for Cancer Nanomedicine. Author contributions: Q.Z., E.K.W.T., and S.N.B. conceived and designed the study. Q.Z., E.K.W.T., C.N., A.S., and T.P. performed experiments. Q.Z., E.K.W.T, and A.S. performed all statistical analysis. Q.Z. generated animal models. Q.Z., E.K.W.T., H.E.F and S.N.B. supervised the research. Q.Z., E.K.W.T., and S.N.B. wrote the first draft of the manuscript. H.E.F. assisted in the preparation of the manuscript. All authors contributed to writing and editing subsequent drafts of the manuscript and approved the final manuscript. We also thank the Comparative Pathology Lab at MIT for clinical chemistry analysis of mouse serum samples, Dr. Liangliang Hao and Dr. Melodi Anahtar from the Bhatia lab for meaningful discussion, Kasia Grzelak and Amy Stoddard for technical assistance, and Prof. Darrell Irvine and Prof. Phillip Sharp at MIT for providing cancer cell lines. Competing interests: S.N.B. reports board members of Brown University, Vertex Pharmaceuticals*, Port Therapeutics*, Ropirio Therapeutics*, and advisory roles in Sunbird Bio*, Satellite Bio*, Matrisome Bio*, Xilio Therapeutics, Danaher, Ochre Bio, Amplifyer Bio*, Earli Inc., Brigham Research Institute/Brigham and Women’s Hospital, and Impilo Therapeutics. *S.N.B. holds equity in these institutions. The remaining authors declare no competing interests. Data and materials availability: All data associated with this study are present in the paper and Supplementary Materials.

## Supplementary figures

**Figure S1**. Optimizing FLIP formulation for optimal membrane fusion.

**Figure S2**. Various molecules are suitable as fusion masks.

**Figure S3**. Cytosolic delivery of small molecules and proteins with C-FLIPs.

**Figure S4**. Synthesis and characterization of C-FLIP/GzmB.

**Figure S5**. Gating of eGFP knockout by C-FLIP/Cas9-gRNA from Hela d2eGFP cells.

**Figure S6**. Protease dysregulation in KP lung tumors.

**Figure S7**. Toxicity evaluation of intravenously injected C-FLIP.

**Figure S8**. Upregulation of co-stimulatory receptors on bone marrow-derived dendritic cells (BMDCs) stimulated by conditional media collected from TC-FLIP/GzmB-treated KP cells.

**Figure S9**. TC-FLIP/GzmB alone significantly suppresses tumor growth and further synergizes with immune checkpoint blockade in a KP lung metastasis model.

**Figure S10**. TC-FLIP/GzmB alone or C-FLIP/GzmB with PD-1 blocking antibodies downregulate cell proliferation biomarker - Ki67.

**Figure S11**. Immunohistochemical staining of gasdermin E expression in healthy lungs and lung metastases.

**Figure S12**. TC-FLIP/GzmB alone suppresses tumor growth and therapeutically synergizes with immune checkpoint blockade in a tumor model of CT26 lung metastasis.

**Figure S13**. TC-FLIP/GzmB alone or with αPD-1 results in significantly higher cleavage of caspase 3 in lung tumors.

**Figure S14**. Phenotypic profiling of T cells in the lungs with tumors over the treatment

