## Supplementary material for "Conditional fusogenic lipid nanocarriers for cytosolic delivery of macromolecular therapeutics": S

### Supplementary information

**Materials.** All chemicals and solvents were purchased from Millipore Sigma (Burlington, MA) unless specified. Lipids:1,2-dioleoyl-3-trimethylammonium-propane (DOTAP), 1,2-dimyristoyl-sn-glycero-3-phosphocholine (14:0 PC, DMPC), 1,2-dimyristoyl-sn-glycero-3-phosphoethanolamine (14:0 PE, DMPE), 1,2-distearoyl-sn-glycero-3-phosphoethanolamine-N-[maleimide(polyethylene glycol)-2000] (DSPe-PEG2k-Mal), 1,2-dioleoyl-sn-glycero-3-phosphoethanolamin [18:1 (Δ9-Cis) PE, DOPE], and 1,2-distearoyl-sn-glycero-3-phosphoethanolamine-N-[methoxy(polyethylene glycol)-2000] (DSPE-mPEG2k) were purchased from Avanti Polar Lipids. DOPE-maleimide (DOPE-Mal) and DSPE-maleimide (DSPE-Mal) were purchased from NOF America Corporation (Boston office, MA). All synthesized peptides purchased from CPC Scientific (Sunnyvale, CA). All nucleic acids unless specified were purchased from Integrated DNA Technologies via custom synthesis (Coralville, IA). KP with or without luciferase transfected cells were obtained from Jacks lab at Massachusetts Institute of Technology (MIT). The cells were isolated from $Kras^{LSL12D}$;$Tp53^{fl/fl}$ lung adenocarcinoma, sorted and transfected with luciferase. CT26 cells transfected with luciferase were attained in our lab. Human umbilical vein endothelial cells (HUVECs). A549 (human lung adenocarcinoma cells), MC38 (mouse colorectal cancer cells), B16F10 (mouse melanoma cells), and J774.2 (mouse macrophages) were obtained from ATCC. DC2.4 (mouse dendritic cells) were obtained by the Irvine lab at MIT. HEK293T cells were obtained from Christopher Chen lab at Boston University. HeLa cells expressing destabilized GFP (HeLa-d2eGFP cells) was kindly provided by Prof. Phillip A. Sharp lab at MIT). Roswell Park Memorial Institute (RPMI) 1640 medium, Dulbecco's Modified Eagle Medium (DMEM), DMEM (high glucose, no glutamine, no phenol red), penicillin/streptomycin solution (P/S; 10,000 units/mL of penicillin, 10,000 µg/mL of streptomycin), L-glutamine (1 M), and β-mercaptoethanol (55mM, 1000X) were purchased from Thermo Fisher Scientific, Granulocyte-macrophage colony-stimulating factor [(GM-CSF) #ab9742)] were purchased from Abcam. EmbryoMax® MEM, non-essential amino acids 100X (#TMS-001-C) was purchased from EMD Millipore. Fetal bovine serum (FBS) was purchased. EnGen Spy Cas9 NLS (#M0646M) was purchased from New England Biolabs (Ipswich, MA).

Antibodies used for immunofluorescent staining on fixed and frozen tissue slides were anti-cytokeratin 8 (CK8), clone TROMA-1 (MABT329) from Millipore Sigma; purified mouse anti-Ki67 (#550609) from BD Biosciences; cleaved caspase-3 (Asp175) antibody (#9661) from Cell Signaling Technology. For antibodies used for flow cytometry, APC mouse anti-MHC Class I (#17-5999-82), APC-eFluor™ 780 mouse anti-MHC Class II (#47-5321-82), FITIC (#11-5773-82) or PE (#12-5773-80) FOXP3 monoclonal antibody (FJK-16s), FITC CD27 monoclonal antibody (LG.7F9) (#11-0271-82), and PerCP-eFluor™ 710 CD127 monoclonal antibody (A7R34) (#46-1271-82) were purchased from Thermo Fisher Scientific (Waltham, MA). Zombie Aqua™ Dye (#77143), FITC anti-mouse CD86 (#105005), APC anti-mouse CD80 (#104713), Brilliant Violet 421™ anti-mouse CD3 (#100227), Brilliant Violet 785™ anti-mouse CD25 (#102051), Brilliant Violet 711™ anti-mouse/human CD44 (#103057), FITC hamster anti-mouse CD40 antibody (#102905), Brilliant Violet 605™ anti-mouse CD45.2 (#109841), PE-Cy7 anti-mouse CD279 (PD-1) antibody (#109109), APC/Fire™ 750 anti-mouse CD62L antibody (#104449), APC anti-mouse CD103 antibody (#121414), PerCP/Cy5.5 anti-mouse CD197 (CCR7) antibody (#120116), and Brilliant Violet 605™ anti-mouse CD366 (Tim-3) antibody (#119721) were purchased from BioLegend (San Diego, CA). Alexa Fluor® 700 mouse anti-Ki-67 (#561277), BUV737 rat anti-mouse CD4 (#612843), BUV737 rat anti-mouse CD86 Clone GL1 (#741437), BD Horizon™ BUV395 rat anti-mouse CD8α (#563786), BD Horizon™ BV510 mouse anti-human granzyme B (#56338), BD OptiBuild™ BV605 hamster anti-rat/mouse CD49a (#740375), and BD Pharmingen™ PE mouse anti-T-bet (#561268) from BD Biosciences (Franklin Lakes, NJ). Alexa

Fluor®647 TCF1/TCF7 (C63D9) rabbit monoclonal antibody (#6709S) was purchased from Cell Signaling Technology (Danvers, MA).

**Preparation of conditional fusogenic liposomal platform (C-FLIP).** The C-FLIP mainly contains phospholipids, synthetic polymers, and synthetic peptides. The lipids used in the C-FLIPs are categorized into cationic, helper, structural and functional lipids with phase transition lower or equal to room temperature. The C-FLIPs were prepared via a traditional extrusion method. A lipid film was formed at predefined lipid ratios (e.g DOTAP/DMPC/DOPE-Mal/DSPE-mPEG2k = 20/70/7/3), and then rehydrated with PBS (1X, pH 7.4). The rehydrated lipids sequentially passed through a mini extruder loaded with a filter membrane of respective 100 nm (10 times back and forth) and 50 nm (11 times back and forth) to form unilamellar FLIPs. Finally, the FLIPs were covalently conjugated with tandem molecules that contain a protease-responsive peptide substrate and a fusion mask. The crude C-FLIPs were purified with via dialysis (MWCO=300kD) against deionized water, size chromatography, or centrifugal decantation. Alternatively, the tandem peptides can also be conjugated to phospholipids before the lipid film preparation, where the conjugation step took place before liposome formation. Similarly, the rehydrated lipids were extruded and purified.

**Conjugation of targeting peptides onto functionalized lipids.** An iRGD peptide with N-terminal cysteine was conjugated to DSPE-PEG2k-Mal at a ratio of thiol/Mal = 1.2/1 in the mixture of dimethyl sulfoxide (DMSO)/chloroform (v/v, 40/60) for overnight at room temperature. The chloroform was fully evaporated and DMSO was exchanged for deionized water using ultrafugal centrifugation. The crude product was then purified with HPLC equipped with a C4 preparatory column (Eluent A=0.05% TFA in $H_2O$, Eluent B=0.043%TFA in acetonitrile/$H_2O$ = 80/20) and with an increasing composition of eluent B (10% to 90%) from 5 min to 70 min. The fractions of interest were collected and lyophilized for further use.

**Conjugation of different fusion masks to C-FLIPs.** Fusion masks tabulated in **Table S1** include short synthetic peptides, zwitterionic molecules, short oligonucleotides, synthetic polymers and antibodies. All these masks are modified to contain thiol groups for a click chemical reaction with maleimide groups from the lipids (e.g., DOPE-MAL, DSPE-MAL) on the FLIPs. Briefly, the overnight reactions were carried out in the pH 7.4 PBS at a ratio of thiol/MAL=1.2/1 at room temperature. The unreacted masks were removed with ultracentrifugal filtration (MWCO=30kD). The resulting C-FLIPs were then stored at 4$^o$C, -20$^o$C or lyophilized for future uses.

**Preparation of control liposomes.** Liposomes formulated as controls comprise phospholipids and PEG-lipids, including LP1 = DOTAP/cholesterol (molar ratio = 50/50), LP2 = DPPC/cholesterol/DSPE-mPEG2k (molar ratio=50/45/5). Similar to C-FLIPs, lipid films of the liposomes were hydrated and liquid extrusion was performed, followed by purification. The lipid compositions of the lipid formulations can be found in **Table S2**. The extrusion of LP2 was performed at 60 $^o$C on a hot plate.

**Freezing and lyophilization of C-FLIPs.** The lipid films prepared as above were rehydrated with 1 mL PBS (1X, pH 7.2-7.4) containing sugar excipients (i.e.,10-50% w/v sucrose, lactose, trehalose or mannitol). The rest of the procedures for synthesizing the C-FLIP were carried out as above. The resulting C-FLIPs were frozen at -80 $^o$C in a “Mr. Frosty” freezing container which guarantees a gradual temperature drop at 1 $^o$C/min. In 2 h, the frozen C-FLIPs were lyophilized in a Labconco FreeZone Benchtop Freeze Dryer. To measure the size, polydispersity and zeta potential, the frozen C-FLIPs were thawed for 0.5 h at room temperature, while the lyophilized C-FLIP powders were redissolved in the same volume of deionized water.

**Confocal microscopy.** Liposomes to be screened were incubated with 50,000 cells per well in a 24-well plate with a glass bottom at a final concentration of lipid 15 or 30 µg/mL for 15 mins. The cells were washed thoroughly with DMEM without phenol red (10% FBS, 1% P/S). The cells were either stained for confocal microscopy or kept cultured in the absence of liposomes for an additional 4 h. The cells were stained with CellBrite® Steady 650 (Biotium #30108-T) at a dilution of 1000X to label the cell membrane, with LysoTracker™ Green DND26 (Invitrogen #L7526) at a dilution factor of 13,334 or LysoTracker™ Deep Red (#L12492) at a dilution factor of 20,000 for acidic compartments such as late endosomes and lysosomes, and followed by Hoechst 33342 (Invitrogen #H3570) staining for nuclei at a dilution factor of 2000. The LysoTracker and Hoechst staining lasted for 45 and 10 mins, respectively. The cells were washed thoroughly with cold PBS for three times, and then incubated in 0.5 mL cold DMEM (10% FBS, 1% P/S) without phenol red for fluorescent or confocal microscopic observation.

**Western blotting.** Cells were treated as indicated for 6 to 24 h and then lysed with RIPA lysis buffer at 100µL/1 million cells with 1X Halt™ phosphate and protease inhibitor cocktail (Thermo Fisher # 78440) for 5 mins on the ice. The cell lysates were gently scraped and then passed 10-15 times through a 25 G needle using a 1ml syringe. The cell lysates were then clarified by centrifugation at 14,000 × g in a microfuge at 4 °C for 10 min. The supernatants were decanted and protein concentrations of different samples were normalized by bicinchoninic acid protein assay (BioRad # 5000001). Protein solutions were denatured at 95 $^oC$ for 10 mins and 20 µg total proteins were loaded per well and beta-actin was used to confirm the similar amount of proteins per well. Cell Signaling Technology antibodies were used to immunoblot for PI3 kinase p85 antibody (#4292), activated (phosphorylated) Akt antibody (#9271), total Akt antibody (#9272), β-actin (8H10D10) mouse mAb (#3700S), caspase-3 antibody (#9662), cleaved caspase-3 (Asp175) antibody (#9661), cleaved gasdermin D (Asp276) rabbit mAb (#10137), gasdermin D (E9S1X) rabbit mAb (#39754), gasdermin E antibody (#40618), and anti-human gasdermin E antibody (#84005).

**Production of recombinant mouse granzyme B.** Double-stranded DNA gBlocks gene fragment encoding the zymogen of mouse granzyme B (GzmB) in **Table S5** that comprises a signal sequence, an enterokinase substrate sequence (replacing native Gly-Glu dipeptide), and a mature granzyme B sequence followed by an octa-histidine tag with flanking SalI and BamHI restriction sites was ordered from Integrated DNA Technologies (Coralville, IA). The gene fragment was cloned into a Genlantis™ gWiz™ vector (Fisher Scientific #P040400) at the SalI and BamHI restriction sites and transformed into DH5α competent *E. coli* cells obtained from New England Biolabs (Ipswich, MA). Selection of the correctly cloned bacterial colony was confirmed by Sanger sequencing (Quintara Biosciences. Cambridge, MA). The plasmid DNA encoding the GzmB zymogen was then isolated from overnight *E. coli* culture using Qiagen Plasmid Midi Kit (Qiagen #12941) following the manufacturer's protocol. GzmB zymogen expression was performed via a transient transfection in FreeStyle™ 293-F cells (Thermo Fisher Scientific # R79007). In brief, the plasmid DNA was pre-incubated with linear polyethylenimine (PEI, 25 kDa, Polysciences # 23966) in OptiPRO™ serum free media (Mass ratio of 1 plasmid: 2 PEI) before adding to a FreeStyle™ 293-F cell suspension culture at $10^6$ cells per mL density and 1 µg/mL final plasmid DNA concentration. After 5 days, the cells were pelleted and the GzmB zymogen in the clarified supernatant was purified via a standard immobilized metal affinity chromatography (IMAC) with Ni-NTA agarose (Qiagen #30210) in Tris buffer. The product was confirmed via SDS-PAGE analysis detected with Coomassie Blue staining. The GzmB zymogen was activated by an overnight incubation with enterokinase from porcine intestine (Sigma #E0885) (1 unit per mg zymogen) at 37°C. The activated GzmB was re-purified with Ni-NTA agarose to remove the enterokinase and buffer-exchanged into PBS using a disposable PD-10 desalting column (Cytiva).

The final product was quantified via 280 nm absorbance, flash-frozen into aliquots, and stored in – 80°C until use.

**Encapsulation of small molecules, peptides, proteins, and ribonucleoprotein particles.** A lipid film was prepared and rehydrated with 0.2 mL sterile PBS (1X, pH 7.2-7.4) containing macromolecular cargos at a concentration of 10 mg/mL of proteins (e.g., Cy5-tagged bovine serum albumin, recombinant mouse granzyme B) or peptides (e.g., peptide-based PROTAC for PI3K in **Table S3**), or of 2 µM of Cas9/sgRNA ribonucleoprotein particles (RNPs; see **Table S4**. The rehydrated lipid vesicles underwent 30-second mild sonication for peptide payloads, or 6 to 10 cycles of freeze-thaw on dry ice and in a 37 $^{o}$C water bath for proteins. The lipid solution was diluted with 0.8 mL of PBS (1X, pH 7.4) and extruded as above. The peptide-, protein or RNP-loaded FLIPs were dialyzed (MWCO=300kD) at 4 $^{o}$C to remove free/unencapsulated cargos. The payloads were quantified with appropriate assays (detailed below). To quantify the drug loading, we first lysed the liposomes with 0.1% Triton X-100 in 1X PBS. The encapsulated peptides were analyzed with Pierce™ Quantitative Colorimetric Peptide Assay (Thermo Fisher Scientific, 23275) per the manufacturer's protocol, which gave 40-45% of encapsulation rate of PROTAC peptides. The loading of proteins (e.g., recombinant granzyme B, bovine serum albumin) was quantified with an SDS-PAGE gel and a standard concentration curve. The FLIP/drug systems were then conjugated with tandem molecules that contain a protease-responsive peptide substrate (e.g., S5, S7, S14 or S28 in Table S3) and a peptide fusion mask (e.g., e8, ekekekekek, DNA10 in **Table S1**) via maleimide-thiol click chemistry to form the C-FLIPs. The conjugation of tandem molecules and subsequent dialysis were found to pose no impacts on the drug loading.

To encapsulate a model small molecule - calcein, a 10 mg/mL solution in dimethylsulfoxide was diluted with PBS at a volume ratio of 10/90. The lipid film was rehydrated with a 0.2 mL of calcein solution and sonicated for 30 s in an ultrasonic bath sonicator at room temperature. The unencapsulated calcein was removed by gel filtration using a Cytiva Sephadex G-100 column. The calcein loading was quantified for fluorescence (Ex/Em = 480/525 nm) on a Tecan M200 Pro microplate reader following the lysis of liposomes with 0.1% Triton X-100.

For Cas9/gRNA RNPs targeting destabilized eGFP of HeLa cells, we first mixed the Cas9 enzyme (New England Biolabs, M0646M) and guide RNA (IDT, custom order) at a molar ratio of 1:1 and incubated the RNP for 5 mins. The RNPs were encapsulated in the FLIP similar to protein encapsulation. The hydrodynamic diameter of the resulting RNPs falls in the range of 15 to 20 nm measured by dynamic light scattering. The RNP loading was measured using a Quant-iT RiboGreen RNA Assay Kit (Thermo Fisher Scientific, R11490) following the lysis of liposomes. As a positive control, the resulting RNP (50 µM Cas9 enzyme) were electroporated into the HeLa expressing destabilized eGFP with a Neon™ NxT Electroporation system (Thermo Fisher Scientific, NEON1SK).

**Cell culture.** Human lung cancer cell lines (HeLa expressing destabilized eGFP) and mouse cancer cell lines (KP, KP expressing luciferase, CT26 expressing luciferase, B16F10, 4T1, MC38, J774A.1, and B16-OVA) were cultured in DMEM supplemented with 10% FBS and 1% penicillin/streptomycin. The media were changed twice every week and the cells were detached for subculture or experiments when they reached approximately 80% confluence.

**Cell death and viability assays**. For assays of survival, 20,000 cells were plated per well in a 96-well plate in 100 µl of medium 24 prior to the assays, with six replicates per condition. The indicated formulations were added in an equal volume of medium and incubated for an additional 24 or 48 h. The medium was then removed and replaced with 100 µl of fresh medium with 20 µL additional CellTiter 96® AQueous One Solution reagent (#G3582 from Progema). The plate was incubated at 37C for 1 h. The absorbance at 490 nm was read on a Tecan M200 Pro microplate reader. Values were normalized to those of PBS-treated control cells. The formulations tested for cell viability are PBS, granzyme b (GzmB), empty FLIP vehicle, non-cleavable FLIP with encapsulated GzmB (N-FLIP/GzmB) or targeting N-FLIP (TN-FLIP/GzmB), and C-FLIP/GzmB pre-cleaved by recombinant MMP9.

We further used a Caspase-Glo® 3/7 Assay System (Promega, G8091) to confirm caspase activity-associated programmed cell death induced by C-FLIP/GzmB following a manufacturer's protocol. Briefly, upon 24 h incubation with the abovementioned samples, we added an equal volume of lysis buffer to cells and monitored the cleavage of peptide substrates by intracellular caspase 3/7 that yielded colorimetric readouts.

**Measurement of immunogenic cell death (ICD) markers.** Calreticulin (CRT), high mobility group box 1 (HMGB1), and adenosine triphosphate (ATP) are well known ICD markers. To measure CRT on the cell surface, 250,000 KP cells per well in a 24-well plate were treated with GzmB, empty FLIP, C-FLIP, non-activated or pre-activated C-FLIP/GzmB for 24 h. All attached and floating cells were harvested and washed twice with the FACS buffer (1X PBS containing 2% heat-inactivated BSA) and incubated with anti-calreticulin antibodies (Thermo Fisher #PA3-900) in presence of anti-Mouse CD16/CD32 (BD Biosciences #553142) for 0.5 h on the ice. The cells were washed once in the FACS buffer and then incubated with goat-anti-rat Alexa Fluor 488-conjugated secondary antibody for 30 mins on the ice, washed twice with the FACS buffer and fixed in 1% paraformaldehyde (PFA) at room temperature for 15 mins. The fixed cells were then resuspended in the FACS buffer, and analyzed by flow cytometry. For HMGB1 measurement in cell culture medium, KP cells were treated for 24 h as described above, medium was collected, and floating cells were removed from the medium by centrifugation at 250 g for 5 mins. Cell-free cell culture media (i.e., conditioned media) were then analyzed by ELISA for HMGB1 (TECAN/IBL-International #30164033) according to the manufacturer's protocol. For measurement of ATP levels, cell-free culture medium obtained as above was analyzed by CellTiter-Glo®2.0 (Promega #G9241) for ATP concentrations according to the manufacturer's protocol. Values were converted to ATP concentrations using a standard curve generated using pure ATP. Interleukin-1β (IL-1 β) in the conditioned media was quantified with an Abcam mouse IL-1 beta ELISA kit (Abcam #ab197742) following the manufacturer's protocol.

**TLR4 activation assay**. HEK-Blue™-mTLR4 reporter cell (InvivoGen #hkb-mtlr4) was cultured a serum-free culture medium supplemented with 50 U/mL penicillin, 50 µg/mL streptomycin, and 100 µg/mL Normocin. A 180 µL of 10,000 HEK-Blue™-mTLR4 cells per well of a 96-well plate were stimulated by additional 20 µL conditioned media of cancer cells treated with different regimens for 24 h. TLR4 activation was revealed colorimetrically with HEK-Blue™ detection assay (InvivoGen #hb-det2). Optical densities were measured with a Tecan M200 Pro microplate reader at 650 nm for absorbance.

**Bone marrow-derived dendritic cells (BMDC) generation.** Bone marrow was harvested from the femurs and tibias of 3 wild-type C57Bl/6 mice (Jackson Laboratories) by flushing out using 1 mL cold RPMI 1640 loaded in a 1 mL syringe and the flushing was repeated once. All the PBS were combined and centrifuged at 4C and 500 g for 5 mins. The cell pellets were then lysed with 1 mL ACK lysis buffer for 3 mins at room temperature to remove red blood corpuscle (RBC) and 10 mL RPMI 1640 were added before the cells were centrifuged again. The resulting cells were filtered through a 100-µm filter to remove aggregates, resuspended at 1 × $10^6$ cells/ml, and cultured in 10-cm non-tissue culture treated petri-dishes (12 million cells per petri-dish) with RPMI 1640 (10% FBS, 1% P/S, 2 mM L-glutamine, 20 ng/ml of GM-CSF, and 55 µM β-mercaptoethanol. After 3 days, 50% of the medium was replaced with fresh medium containing growth factors. Bone marrow-derived dendritic cells (BMDCs), which were suspended or loosely adherent, were harvested by gentle pipetting on the Day 6 (the day for bone marrow harvesting was considered Day 0), and were used for the assay.

***In vitro* cross-presentation assay.** B16-OVA cells were treated with GzmB, empty FLIP and C-FLIP, non-activated and pre-activated C-FLIP/GzmB for 24 h, followed by extensive wash with DMEM (10% FBS, 1% P/S). Subsequently,1 × $10^6$ treated cells were co-cultured with BMDCs per well at a ratio of 4:1. After 24 h of co-culture, supernatants were removed from each well, and the BMDCs were washed two to three times in T cell medium (RPMI 1640 containing 10% FBS, 1% P/S, 20 mM HEPES, 1 mM sodium pyruvate, 55 μM β-mercaptoethanol, 2 mM l-glutamine, and non-essential amino acids). CD8+ OT-1 T cells isolated from spleens of OT-1 mice using EeasySep mouse CD8+ T cell isolation kit (Stemcell Technologies) were then co-cultured with the BMDCs at 125,000 T cells per well and a T cell–to–BMDC ratio of 1:2. After a 14-h incubation, IFN-γ–producing T cells were identified and quantified by intracellular cytokine staining and flow cytometry using a LSR Fortessa flow cytometer. Cells were first gated for CD3 expression and then gated for CD8 and IFN-γ expression among the CD3+ population.

**Pharmacokinetics and systemic biodistribution**. DiR-labeled liposomal formulations including FLIPs and C-FLIPs were injected into wild-type C57Bl/6 mice (n=3 per cohort) via tail veins in a 100 µL sterile PBS containing 1.5 mg/mL total lipids. Blood samples were retro-orbitally collected with heparinized capillary tubes at different timepoints. The 40 µL blood from each mouse was mixed with 40 µL heparin solution (10U/mL), and the mixture was centrifuged at 1000 x g at 4 $^{o}$C for 5 mins. Thirty µL plasma (including heparin solutions) — the liquid portion of blood, was introduced to every well of a 384-well plate measured for DiR fluorescence with a LI-COR Odyssey® DLx imager (LI-COR Biosciences). The first blood was sampled at 1 minute post i.v. injection of the liposomes to which all other samples were normalized as percentages (i.e., DiR fluorescence signal intensity at 1 minute post injection was set 100%). For the systemic biodistribution study, the mice dosed with liposomal formulations were euthanized at 24 h post-injection. The tissues of lung, heart, liver, spleen, and kidney were harvested and imaged for DiR fluorescence using a LI-COR Odyssey® DLx imager.

**Lung metastasis models in mice.** Half a million of luciferase-expressing KP lung tumor cells in 0.1 mL cold sterile 1X PBS were intravenously injected via tail veins into C57BL/6 mice. We used *in vivo* imaging system (IVIS) to monitor tumor proliferation every 4 days starting at Day 3 post tumor cell inoculation. The tumors were allowed to grow until *in vivo* efficacy studies or other assays.

**Lung homogenization**. Healthy lungs and tumor-bearing lungs were harvested and chopped into small pieces using surgical scissors. The tissues were added into pre-chilled 1X PBS at a mass concentration of 100 mg/mL and then placed in GentleMACS C-tubes on the ice. The lung tissues were then dissociated with a GentleMACS dissociator. Centrifuge at 2000 rpm and 4C for 10 min to collect the supernatant. The proteins in the supernatant were quantified using a Bradford assay (Bio-Rad Protein Assay Kit I 5000001) and diluted to 0.5 mg protein/mL for peptide substrate screening.

***In vitro* screening of substrate peptides against lung homogenates.** Fluorogenic protease substrates were synthesized by CPC Scientific customer (Table S3). The fluorogenic substrates P1-P37 (60 μM final concentration) were incubated in 30-μl final volume in 1X PBS with 5 µL homogenate supernatant at 37°C. Proteolytic cleavage of substrates was quantified by increase in fluorescence over time on a Tecan Infinite M200 Pro plate reader. Initial enzyme cleavage rates were calculated via dividing the fluorescence intensity by elapsed time. The fold change of initial cleavage rate of tumor-bearing lung homogenates to that of healthy lungs was used to characterize their protease activity.

***In vivo* toxicity assessment**. For *in vivo* biosafety validation, healthy wild-type C57B6/L mice were intravenously injected with FLIPs and MMP9-cleavable C-FLIPs at a total lipid 15 mg/kg for 5 doses with dosing frequency every other day. Twenty-four h after the last dose, approximately 0.6-0.8 mL blood were collected via cardiac puncture from the mice under deep isoflurane anesthesia. The blood samples were introduced into microtubes with clotting activator gels, and centrifuged at 10,000 x g for 10 mins at 4 $^{o}$C. At least 250 µL serum per mouse were used for blood clinical chemistry tests by the Comparative Pathology Laboratories at Massachusetts Institute of Technology. A panel of 25 analytes was examined, which are ALK phosphatase, ALT, AST, CK, GGT, albumin, total protein-chem, globulin, total bilirubin, BUN, creatinine, cholesterol, glucose, calcium, phosphorus, $TCO_2$ bicarbonate, chloride, potassium, sodium, A/G ratio, B/C ratio, indirect bilirubin, Na/K ratio, and anion gap. For general histopathological evaluation, brain, heart, lungs, liver, kidneys, and spleen were subsequently excised and fixed immediately in 4% paraformaldehyde (PFA) PBS. The fixed tissues were paraffinized and sectioned for hematoxylin and eosin (H&E) staining by the Hope Babette Tang (1983) Histology Facility in the Koch Institute of Integrative Cancer Research, Massachusetts Institute of Technology. The stained slides were histopathologically evaluated by a highly experienced veterinary pathologist, Dr. Roderick Bronson from Harvard Medical School.

**Immunogenicity evaluation**. A 200 µL of 30 mg/kg liposomes in PBS was injected via tail veins. After 24 h, approximately 0.2 mL blood (6 to 7 drops) were sampled into microtubes with clotting activator gels via cheek bleeding. The samples were allowed to stand for 30 mins at the room temperature, and centrifuged at 10,000 x g for 10 mins at 4 $^{o}$C. Fifty µL serum was diluted with an equal volume of sterile PBS (1X, pH 7.4). The serum sample were examined for a panel of 32 cytokines and chemokines (Eve Technologies, Canada) using a Luminex-based multiplex assay, which are eotaxin, G-CSF, GM-CSF, IFNγ, IL-1α, IL-1β, IL-2, IL-3, IL-4, IL-5, IL-6, IL-7, IL-9, IL-10, IL-12p40, IL-12p70, IL-13, IL-15, IL-17A, IP-10, KC, LIF, LIX, MCP-1, M-CSF, MIG, MIP-1α, MIP-1β, MIP-2, RANTES, TNFα, and VEGF-A.

***In vivo* antitumor experiment**. All animal studies and procedures were approved by the Committee for Animal Care at Massachusetts Institute of Technology. Half million KP cells transfected luciferase were injected into age-matched female or male wild-type C57BL/6 mice via the tail veil and the mice were randomly regrouped into 5 mice per cohort. Seven days after the injection of tumor cells, therapeutic formulations were intravenously administered at a pre-determined amount for 3 doses a week and 5 doses in total. Tumor burdens in the KP tumor-bearing lung was longitudinally monitored with an *in vivo* imaging system for luciferase luminescence until an indicated terminal date or a total luminescence signal of tumors reaching $5X10^9$ where the mice were euthanized). Different regimens given to the mice were i) i.v. PBS, ii) i.v. GzmB 3 mg/kg per dose, iii) i.v. TN-FLIP/GzmB (targeting, non-cleavable FLIP with GzmB encapsulated) at 3 mg/kg GzmB, iv) i.v. MMP9-activatable C-FLIP$^{S7}$/GzmB at 3 mg/kg GzmB, v) i.p. αPD-1 (immune checkpoint PD-1 antibody) at 10mg/kg per dose, and vi) i.v. C-FLIP/GzmB (3mg/kg per dose) in combination with i.p. αPD-1 (10 mg/kg per dose), viii) i.v. LNP/SiR7 (siR equivalent 100 µg/kg), and ix) i.v. LNP/SiR7 (siR equivalent 100 µg/kg) in combination with i.p. αPD-1 (10 mg/kg per dose). At the terminal point, the lungs were excised and fixed with 4% PFA for 24 h. H&E, immunohistochemical, or immunofluorescent staining were carried out on the fixed lung tissues for cleaved caspase 3, Antigen Kiel 67 (Ki67), cytokeratin 8 (CK8), or gasdermin E. For the lung metastasis of colorectal cancer - CT26 with luciferase transfected, 200k cells were intravenously injected, treated and monitored for tumor burdens as above.

To assess the memory T cell populations upon combinatorial treatment of TC-FLIP/GzmB and αPD-1, 200,000 luciferase-transfected KP cells were injected into age-matched female or male wild-type C57Bl/6 mice via the tail veil and the mice were randomly regrouped into 10 mice per cohort. Five days after the injection of tumor cells, TC-FLIP/GzmB and αPD-1 were intravenously administered at a pre-determined amount for 3 doses a week and 6 doses in total. The Bruker Skyscan 1276 microCT imaging system was used to monitor the dynamics of lung tumors along the treatment. At the terminal point (i.e., a week post last treatment or when meets endpoint criteria), complete response (CR) is defined as no detectable lung nodules with microCT, the tumors shrinking (>30%) but not disappearing during the observation are grouped into partial response (PR), while delayed tumor growth without shrinking is defined as progressive diseases (PD). At Day 23 (one week post last treatment), the lungs were harvested to prepare for single cell suspension as below for profiling lung-resident memory T cell populations using flow cytometry.

**Flow cytometry**. To assess the activation of dendritic cells (DCs) by the conditioned media isolated from the KP cells treated with different liposomal regimens, 200,000 BMDCs were seeded per well in a non-tissue culture treated, round-bottom 96-well plate and incubated in 180 µL R10 medium and 20 µL conditioned media (i.e., total volume 200 µL). In 24 h, the cells were harvested and washed with FACS buffer (2% heat-inactivated FBS in PBS). The cells were blocked by anti-mouse CD16/CD32, and then stained with Zombie Aqua™ Dye (#77143), BUV737 anti-mouse CD86, APC anti-mouse CD80, FITC anti-mouse CD40, APC-eFluor™ 780 mouse anti-MHC Class II, and APC mouse anti-MHC Class I in presence of the CD16/CD32 antibody. The stained BMDCs were fixed with 1% PFA in PBS for 15 mins at the room temperature. the activation markers in the DCs were quantified with flow cytometry using an LSR Fortessa flow cytometer and analyzed with the FlowJo software.

At the terminal point of the mice treated with various regimens, to distinguish tissue-resident (i.e., lung, mDLNs, and spleen) versus circulating immune cells, mice were anesthetized and injected retro-orbitally with a fluorescently-conjugated anti-CD45 antibody (PE-CF594 or AlexaFluor780; 30-F11; BD Bioscience) 2 mins prior to euthanasia. Circulating CD8 and CD4 T cells stained positive for the CD45 antibody were excluded from our analyses. After 2 mins, the mice were immediately euthanized with the excess of isofluorane. The spleen, mediastinal draining lymph nodes (mDLNs), and lung tissues were collected and carefully minced by hand with spring scissors and incubated in 125 U/mL collagenase IV (Worthington Biochemical) and 40 U/mL Roche DNase I (Sigma-Aldrich, 04536282001) for 30 mins at 37°C. Following incubation, the tissue from every mouse was passed through a 100 µm cell strainer. In the meantime, spleen and mDLNs were dissociated with a plunger (rubber end) of a BD 1 mL syringe and then passed through a 70 µm cell strainer into RPMI 1640 media containing 2% heat-inactivated FBS. Cell suspensions were pelleted by centrifugation (300 RCF for 5 mins at 4 $^{o}$C), resuspended in 1 mL 1X ACK lysis buffer and incubated on ice for 2 mins to lyse red blood cells. Cells were pelleted again and resuspended in 1X PBS supplemented with 2% heat-inactivated FBS. The single cell suspensions were transferred to a 96-well U-bottom plate (non-tissue culture treated). Cells were then stained with a fixable viability dye to exclude dead cells (1:200 dilution; 20 mins on ice; Zombie Aqua™ Dye, BioLegend #77143 or Tonbo Ghost red 710 Dye, Tonbo Biosciences #13-0871) and resuspended in the FACS buffer (2% heat-inactivated FBS in 1X PBS) and stained with a following panel (Panel 1) of surface antibodies in presence of CD16/CD32 Fc antibody (BD Biosciences #553142) for 30 mins on ice: BV421 CD3 (BioLegend #100227), BUV395 CD8α (BD Biosciences #563786), BUV737 CD4 (BD Biosciences, #612843), BV785 CD25 (BioLegend #102051), BV 711™ CD44 (BioLegend #103057), PerCP-Cy5.5 CD69 (Invitrogen #45-0691-80), PE-Cy7 CD279 (i.e., PD-1) (BioLegend #109109), and BV605™ CD366 (i.e., Tim-3) (BioLegend #119721). For intracellular staining of transcriptional factors and cytoplasmic proteins, the cells were fixed for overnight (~12-16 h) at 4 $^{o}$C using the eBioscience™ Fixation/Permeabilization buffer (Invitrogen #00-5123-43), followed by 20-minute permeabilization with 1X working solution of Permeabilization Buffer (Invitrogen #88-8824-00). Next, the cells were then stained for 1 h at the room temperature with the following antibodies in presence of 1X permeabilization buffer: eBioscience™ PE Foxp3 (Invitrogen #12-5773-80), Alexa Fluor®647 TCF1/TCF7 (C63D9) (Cell Signaling Technology #6709S), Alexa Fluor® 700 Ki67 (BD Biosciences #561277), and BV510 granzyme B (BD Biosciences #563388).

In another cohort of tissue samples, the cells were stained with another panel (Panel 2) of surface antibodies for 30 mins on ice: BV421 CD3 (BioLegend #100227), BUV395 CD8α (BD Biosciences #563786), BUV737 CD4 (BD Biosciences, #612843), FITC CD27 (Invitrogen #11-0271-82), BV605 CD49a (BD Biosciences #740375), BV711 CD44 (BioLegend #103057), PerCP-Cy5.5 CD69 (Invitrogen #45-0691-80), APC/Fire 750 CD62L (BioLegend #104449), APC CD103 (BioLegend #121414), PerCP-eFluor 710 CD127 (Invitrogen #46-1271-82), and PerCP/Cy5.5 CD197 (CCR7) (#120116). For intracellular staining, the cells were stained for 1 h at the room temperature with the following antibodies: Alexa Fluor® 700 Ki67 (BD Biosciences #561277) and PE T-bet (BD Biosciences #561268). Samples were analyzed on a BD Biosciences LSR Fortessa I Flow Cytometry Analyzer.

**Table S1. Sequences of fusion masks**

| Category of fusion mask | Annotation | Fusion mask# |
|---|---|---|
| Peptides | $e_8$ | eeeeeeee |
| | $(ek)_5$ | ekekekekek |
| | iRGD | CGGCRGDKGPDC (C2&C3 bridge) |
| Synthetic polymer | PEG2k | polyethylene glycol 2000 |
| Nucleic acid | DNA | T*C*A*T*A*G*T*T*A*G* |

### Peptide sequence indicated from N to C terminus, where lower case letter represents a D-isomer amino acid. denotes d-amino acids. DNA sequence indicated from 5' to 3', where * denotes phosphorothioate (PS) backbone.

**Table S2. Lipid components of FLIPs and C-FLIPs**

| Liposome | Lipid composition | Molar ratio | Fusion mask | Peptide substrate |
|---|---|---|---|---|
| LP1 | DOTAP/cholesterol | 50/50 | - | - |
| LP2 | DPPC/cholesterol/DSPE-mPEG2k | 50/45/5 | - | - |
| FLIP | DOTAP/DMPC/DOPE-MAL/DSPE-PEG-MAL | 20/70/7/3 | - | - |
| $\text{C-FLIP}_{e8}^{S5}$ | DOTAP/DMPC/DOPE-MAL/DSPE-PEG-MAL | 20/70/7/3 | eeeeeeeee | GGPVPLSLVMGC |
| $\text{C-FLIP}_{e8}^{S7}$ | DOTAP/DMPC/DOPE-MAL/DSPE-PEG-MAL | 20/70/7/3 | eeeeeeeee | GGPLGVRGKGC |
| $\text{C-FLIP}_{e8}^{S14}$ | DOTAP/DMPC/DOPE-MAL/DSPE-PEG-MAL | 20/70/7/3 | eeeeeeeee | GGGSGRSANAKGGC |
| $\text{C-FLIP}_{e8}^{S28}$ | DOTAP/DMPC/DOPE-MAL/DSPE-PEG-MAL | 20/70/7/3 | eeeeeeeee | Nle(Obzl)-M(O)2-Oic-Abu-C |
| $\text{N-FLIP}_{e8}^{None}$ | DOTAP/DMPC/DOPE-MAL/DSPE-PEG-MAL | 20/70/7/3 | eeeeeeeee | N/A |

**Table S3. Peptide sequences.**

| Peptide substrate | Sequence |
|---|---|
| GzmB substrate | 5FAM-GGAIEFDSG-K(CPQ2)-PEG2-C |
| P1 | 5FAM-GGPQGIWGQK(CPQ2)-PEG2-C |
| P2 | 5FAM-GGLVPRGSGK(CPQ2)-PEG2-C |
| P3 | 5FAM-GGPVGLIGK(CPQ2)-PEG2-C |
| P4 | 5FAM-GGPWGIWGQGK(CPQ2)-PEG2-C |
| P5 | 5FAM-GGPVPLSLVMK(CPQ2)-PEG2-C |
| P6 | 5FAM-GGPLGLRSWK(CPQ2)-PEG2-C |
| P7 | 5FAM-GGPLGVRGKK(CPQ2)-PEG2-C |
| P8 | 5FAM-GGf-Pip-RSGGGK(CPQ2)-PEG2-C |
| P9 | 5FAM-GGfPRSGGGK(CPQ2)-PEG2-C |
| P10 | 5FAM-GGf-Pip-KSGGGK(CPQ2)-PEG2-C |
| P11 | 5FAM-GRQRRALEKG-K(CPQ2)-PEG2-GC |
| P12 | 5FAM-GILSRIVGGG-K(CPQ2)-PEG2-GC |
| P13 | 5FAM-GKPISLISSG-K(CPQ2)-PEG2-GC |
| P14 | 5FAM-GGGSGRSANAKG-K(CPQ2)-PEG2-GC |
| P15 | 5FAM-GGGPG-K(CPQ2)-PEG2-GC |
| P16 | 5FAM-GRPKPVE(Nval)WRKG-K(CPQ2)-PEG2-GC |
| P17 | 5FAM-GHSSKLQG-K(CPQ2)-PEG2-GC |
| P18 | 5FAM-GSSQYSSNGG-K(CPQ2)-PEG2-GC |
| P19 | 5FAM-GQKGRYKQEG-K(CPQ2)-PEG2-GC |
| P20 | 5FAM-GGKAFRRSGG-K(CPQ2)-PEG2-GC |
| P21 | 5FAM-GIQQRSLGGG-K(CPQ2)-PEG2-GC |
| P22 | 5FAM-GGVPRGG-K(CPQ2)-PEG2-GC |
| P23 | 5FAM-GSGSKIIGGG-K(CPQ2)-PEG2-GC |
| P24 | 5FAM-GAANLTRG-K(CPQ2)-PEG2-GC |
| P25 | 5FAM-GGGELRG-K(CPQ2)-PEG2-GC |
| P26 | 5FAM-GLAQAPhe(homo)RSG-K(CPQ2)-PEG2-GC |
| P27 | 5FAM-GSPLAQAVRSSG-K(CPQ2)-PEG2-GC |
| P28 | 5FAM-Nle(Obzl)-M(O)2-Oic-Abu-K(CPQ2)-PEG2-C |
| P29 | 5FAM-GPVPLSLVMG-K(CPQ2)-PEG2-GC |
| P30 | 5FAM-GRQSRIVGGG-K(CPQ2)-PEG2-GC |
| P31 | 5FAM-GSQPRIVGGG-K(CPQ2)-PEG2-GC |
| P32 | 5FAM-GPLGMRGKGG-K(CPQ2)-PEG2-GC |
| P33 | 5FAM-GAPRWIQDKGG-K(CPQ2)-PEG2-GC |
| P34 | 5FAM-GRPPGFSAFKGG-K(CPQ2)-PEG2-GC |
| P35 | 5FAM-GSYRIFGG-K(CPQ2)-PEG2-GC |
| P36 | 5FAM-RPKPVEGG-K(CPQ2)-PEG2-GC |
| P37 | 5FAM-GRRRGGAANC(OMeBzl)RMGG-K(CPQ2)-PEG2-GC |

| | |
|---|---|
| S1 | 5FAM-GGPQGIWGQGC |
| S2 | 5FAM-GGLVPRGSGGC |
| S3 | 5FAM-GGPVGLIGGC |
| S4 | 5FAM-GGPWGIWGQGC |
| S5 | 5FAM-GGPVPLSLVMGC |
| S6 | 5FAM-GGPLGLRSWGC |
| S7 | 5FAM-GGPLGVRGKGC |
| S8 | 5FAM-GGf-Pip-RSGGGC |
| S9 | 5FAM-GGfPRSGGGC |
| S10 | 5FAM-GGf-Pip-KSGGC |
| S11 | 5FAM-GRQRRALEKGC |
| S12 | 5FAM-GILSRIVGGGC |
| S13 | 5FAM-GKPISLISSGGC |
| S14 | 5FAM-GGGSGRSANAKGGC |
| S15 | 5FAM-GGGPGGC |
| S16 | 5FAM-GRPKPVE(Nval)WRKGGC |
| S17 | 5FAM-GHSSKLQGGC |
| S18 | 5FAM-GSSQYSSNGGC |
| S19 | 5FAM-GQKGRYKQEGGC |
| S20 | 5FAM-GGKAFRRSGGGC |
| S21 | 5FAM-GIQQRSLGGGC |
| S22 | 5FAM-GGVPRGGGC |
| S23 | 5FAM-GSGSKIIGGGC |
| S24 | 5FAM-GAANLTRGGC |
| S25 | 5FAM-GGGELRGGC |
| S26 | 5FAM-GLAQAPhe(homo)RSGGC |
| S27 | 5FAM-GSPLAQAVRSSGGC |
| S28 | 5FAM- Nle(Obzl)-M(O)2-Oic-Abu-C |
| S29 | 5FAM-GPVPLSLVMGGC |
| S30 | 5FAM-GRQSRIVGGGC |
| S31 | 5FAM-GSQPRIVGGGC |
| S32 | 5FAM-GPLGMRGKGGGC |
| S33 | 5FAM-GAPRWIQDKGGGC |
| S34 | 5FAM-GRPPGFSAFKGGGC |
| S35 | 5FAM-GSYRIFGGGC |
| S36 | 5FAM-RPKPVEGGGC |
| S37 | 5FAM-GRRRGGAANC(OMeBzl)RMGGC |
| pP85 | NH2-GPGGD(pY)AAMGACPASEQG(pY)EEMRA-PEG3-ALAPYIP-CONH2* |
| CPP-pP85 | NH2-GPGGD(pY)AAMGACPASEQG(pY)EEMRA-PEG3-ALAPYIP-(rrrrrrrr)-CONH2* |

* pY denotes phosphorylated tyrosine.

##### Table S4. DNA or RNA sequences

| | |
|---|---|
| **Custom d2eGFP sequence for Hela d2eGFP cells** | ATGGTGAGCAAGGGCGAGGAGCTGTTCACCGGGGTGGTGCCCATCCTG<br>GTCGAGCTGGACGGCGACGTAAACGGCCACAAGTTCAGCGTGTCCGGC<br>GAGGGCGAGGGCGATGCCACCTACGGCAAGCTGACCCTGAAGTTCATC<br>TGCACCACCGGCAAGCTGCCCGTGCCCTGGCCCACCCTCGTGACCACC<br>CTGACCTACGGCGTGCAGTGCTTCAGCCGCTACCCCGACCACATGAAG<br>CAGCACGACTTCTTCAAGTCCGCCATGCCCGAAGGCTACGTCCAGGAG<br>CGCACCATCTTCTTCAAGGACGACGGCAACTACAAGACCCGCGCCGAG<br>GTGAAGTTCGAGGGCGACACCCTGGTGAACCGCATCGAGCTGAAGGGC<br>ATCGACTTCAAGGAGGACGGCAACATCCTGGGGCACAAGCTGGAGTAC<br>AACTACAACAGCCACAACGTCTATATCATGGCCGACAAGCAGAAGAACG<br>GCATCAAGGTGAACTTCAAGATCCGCCACAACATCGAGGACGGCAGCG<br>TGCAGCTCGCCGACCACTACCAGCAGAACACCCCCATCGGCGACGGCC<br>CCGTGCTGCTGCCCGACAACCACTACCTGAGCACCCAGTCCGCCCTGA<br>GCAAAGACCCCAACGAGAAGCGCGATCACATGGTCCTGCTGGAGTTCG<br>TGACCGCCGCCGGGATCACTCTCGGCATGGACGAGCTGTACAAGAAGC<br>TTAGCCATGGCTTCCCGCCGGAGGTGGAGGAGCAGGATGATGGCACGC<br>TGCCCATGTCTTGTGCCCAGGAGAGCGGGATGGACCGTCACCCTGCAG<br>CCTGTGCTTCTGCTAGGATCAATGTGTAG |
| **Target DNA sequence** (antisense strand) | GGGCACGGGCAGCTTGCCGG |
| Ordered chemically synthetic sgRNA sequence for Hela d2eGFP cells: | 5'**GGGCACGGGCAGCUUGCCGG**GUUUUAGAGCUAGAAAUAGCAAGUUA<br>AAAUAAGGCUAGUCCGUUAUCAACUUGAAAAAGUGGCACCGAGUCGG<br>UGCUUUUUU3' |

##### Table S5. Amino acid and corresponding gBlock sequences of recombinant mouse pro-granzyme B.

**Amino acid sequence:**

MKILLLLLTLSLASRTKADDDDKIIGGHEVKPHSRPYMALLSIKDQQPEAICGGFLIREDFVLTAAHCEGS
IINVTLGAHNIKEQEKTQQVIPMVKCIPHPDYNPKTFSNDIMLLKLKSKAKRTRAVRPLNLPRRNVNVKP
GDVCYVAGWGRMAPMGKYSNTLQEVELTVQKDRECESYFKNRYNKTNQICAGDPKTKRASFRGDS
GGPLVCKKVAAGIVSYGYKDGSPPRAFTKVSSFLSWIKKTMKSSASHHHHHHHH

**gBlock sequence:**
ATTATCGGAGGTCACGAAGTGAAACCACATTCCCGACCGTATATGGCATTGCTTTCCATAAAGGAT
CAGCAACCGGAGGCCATATGTGGGGGCTTCCTTATCCGCGAGGACTTTGTTCTTACCGCTGCCCA
TTGTGAGGGTTCAATTATTAACGTGACGTTGGGGGCGCATAATATCAAAGAACAGGAGAAGACGC
AACAAGTAATCCCGATGGTGAAGTGCATTCCGCATCCAGATTACAATCCTAAGACCTTCAGTAATG
ACATTATGCTCCTTAAACTTAAGTCAAAGGCGAAACGAACACGAGCAGTACGGCCCCTTAACCTGC
CGCGAAGGAATGTAAACGTGAAGCCCGGTGACGTCTGTTATGTGGCTGGATGGGGAAGGATGGC
ACCCATGGGTAAATATTCAAACACTCTCCAGGAGGTGGAACTGACCGTACAAAAAGACCGCGAAT
GCGAATCTTATTTCAAAAACAGATATAATAAGACAAACCAAATCTGTGCGGGAGACCCAAAGACAA
AGAGAGCAAGTTTTAGGGGAGATTCTGGCGGCCCACTTGTTTGCAAGAAGGTGGCCGCAGGGAT
TGTGTCATACGGCTACAAGGATGGATCACCCCCTCGCGCATTCACCAAGGTAAGCTCATTTCTGA
GTTGGATTAAAAAGACTATGAAGTCCTCTTGA

**Table S6. Primer sequences for amplifying gBlock of recombinant granzyme B and**

| Primer name | Forward or reverse | Sequence |
|---|---|---|
| PAT5 | Forward | AGACATAATAGCTGACAGAC |
| PAT13 | Forward | CCTTTCCATGGGTCTTTTCTGCAGTCACCGTCGTCGACACCGG TGCCGC |
| PAT11 | Reverse | GGCAACTAGAAGGCACAG |
| CN230 | Reverse | CTAGAGCTCTGATCTTTTATTAGCCAGAAGTGATCTGGATCCT CAATGATGGTGATG |

A

| Formulation | DOTAP | DOPE-MAL | DSPE-PEG2k | DMPC | Size (d.nm) | Polydispersity index (PDI) | Zeta potential (mV) |
|---|---|---|---|---|---|---|---|
| FLIP_DOTAP10 | 10 | 7 | 3 | 80 | 89.7±0.7 | 0.04±0.02 | 0.21±0.1 |
| FLIP_DOTAP20 | 20 | 7 | 3 | 70 | 89.7±0.5 | 0.06±0.01 | 14.4±0.2 |
| FLIP_DOTAP50 | 50 | 7 | 3 | 40 | 92.1±1.1 | 0.04±0.04 | 15.8±0.9 |

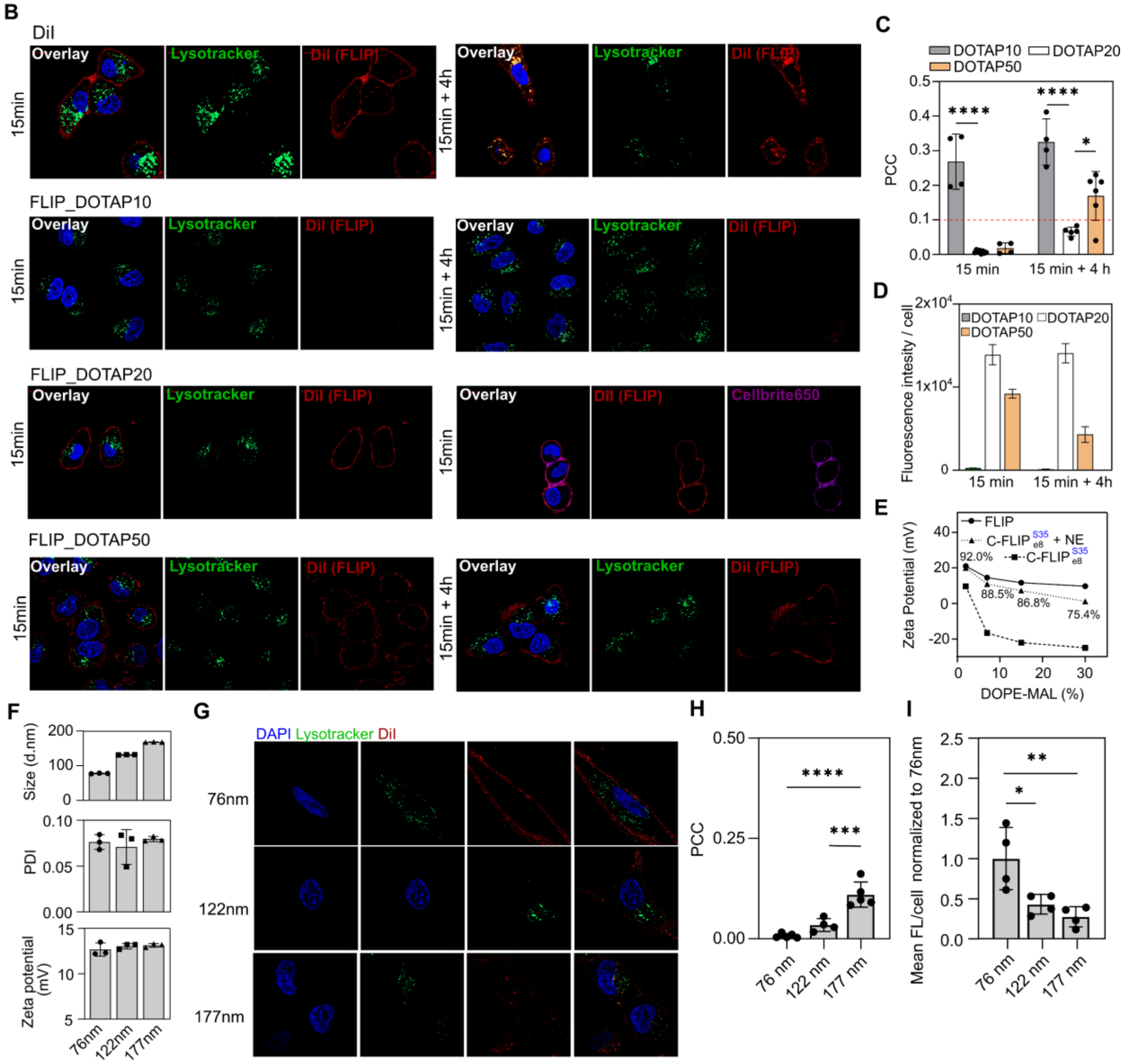


**Figure S1**. **Optimizing FLIP formulation for optimal membrane fusion.** Cationic lipids dictate FLIP's ability to fuse with cell membrane. **(A)** Size, polydispersity index (PDI) and zeta potential of FLIPs with different DOTAP ratios. Size and PDI remain unchanged, whereas surface charges are from moderately positive to nearly neutral as the DOTAP ratio declines from 50% to 10%. FLIP_DOTAP10, 20, 50 denotes that the molar ratio of DOTAP among all lipid components is 10, 20 and 50%, respectively. **(B)** Lipophilic dye DiI and liposomes (FLIP_DOTAP10, 20, and 50) were incubated with mouse KP lung cancer cells for 15 mins before being washed away and were live imaged immediately (left column) or 4 h after wash to monitor membrane fusion (right column) as benchmarked against 15 mins. Lipophilic DiI quickly labels both cellular plasma membranes and endolysosomal membranes (within 15 min incubation) and its accumulation in endosomes and lysosomes increases over time (i.e., 15min+4h). FLIP_DOTAP10 has negligible fusion with the cellular membrane in comparison with those with FLIP_DOTAP20 and FLIP_DOTAP50. **(C)** Pearson colocalization coefficient (PCC) confirms FLIP_DOTAP10 is highly colocalized with endosomes and lysosomes, whereas FLIP_DOTAP20 and FLIP_DOTAP50 demonstrate high and moderate fusion with plasma membrane, respectively. FLIP_DOTAP20 is thus utilized in the following experiments to construct conditional fusogenic liposomes (C-FLIPs). **(D)** FLIPs with higher DOTAP ratios tend to fuse and/or be taken up by cells as indicated by overall fluorescence intensity per cell. **(E)** Restoration of surface charge of C-FLIP is inversely proportional to the percentage of functional DOPE-MAL in the lipid components. **(F)** Size, PDI and zeta potential of the FLIP_DOTAP20 with different sizes (i.e., 76, 122 and 177 nm). The surface charge of the FLIPs largely remains the same, with varying size. **(G)** Confocal microscopy and **(H)** colocalization analysis reveal that the FLIP with a smaller size (76 nm) tends to effectively fuse with cell membrane, while the increased portions of larger FLIPs are endocytosed. **(I)** Particle-membrane interaction (including adsorption, fusion and endocytosis) decreases when FLIP size increases.

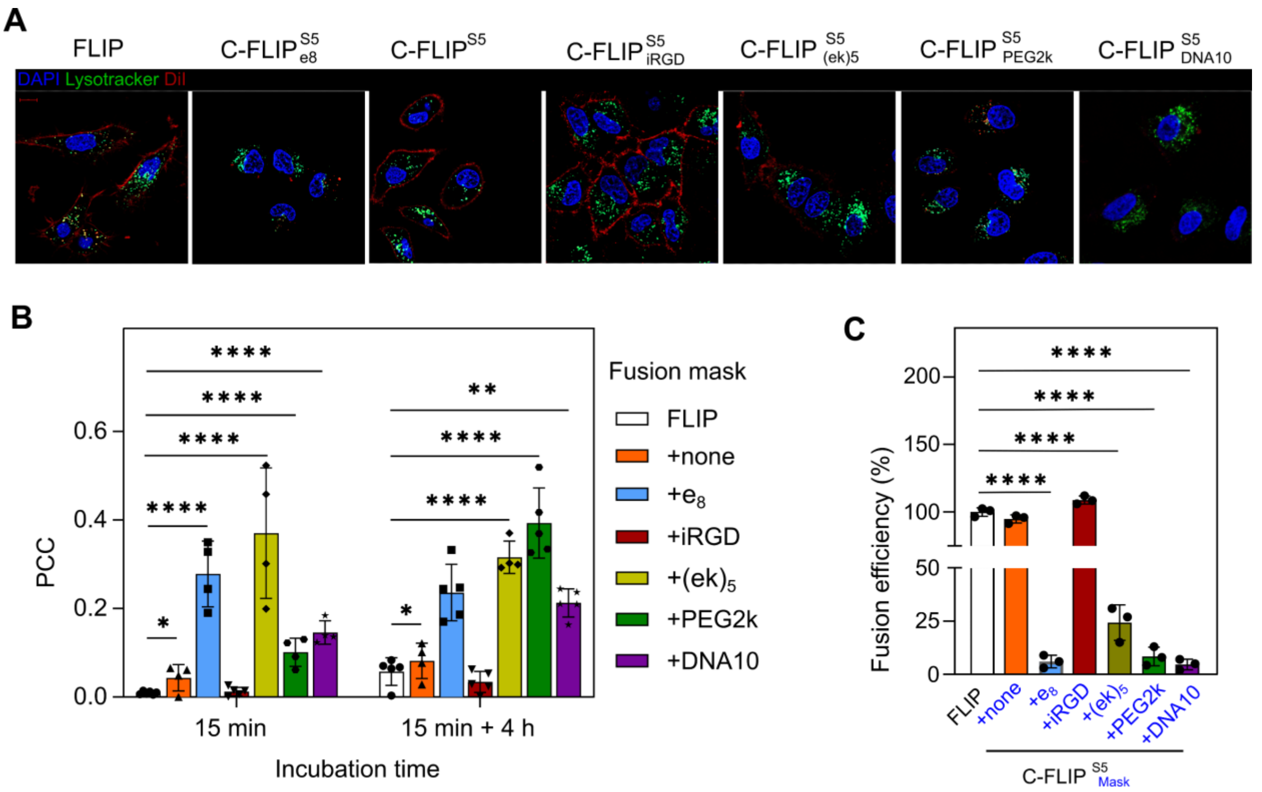


**Figure S2. Various molecules are suitable as fusion masks. (A)** Effects of various motifs on the fusogenicity of the liposomes. The molecules (denoted in subscripts of) were conjugated to the FLIP_DOTAP20 via a peptide S5, a MMP substrate, as indicated in superscripts (C-$FLIP^{S5}_{fusion\ mask}$). The examined motifs are peptides ($e_8$, iRGD, $(ek)_5$), polyethylene glycol 2000Da (PEG2k), and single stranded oligonucleotide with 10 base pairs (DNA10). The detailed sequences can be referred to **Table S1. (B)** Pearson colocalization coefficient (PCC) of various C-FLIPs with the lysotrackers. The C-FLIPs are conjugated with the indicated molecules in (A). Liposomes were incubated with the mouse KP lung cancer cells for 15 mins before they were thoroughly washed. The cells were imaged immediately (i.e., 15 min) after washes. **(C)** Fusion blocking efficiency of indicated motifs upon the normalization to that of FLIP which is designated 100% fusion. The lower fusion efficiency, the higher likelihood of fusion blockade by the motifs.

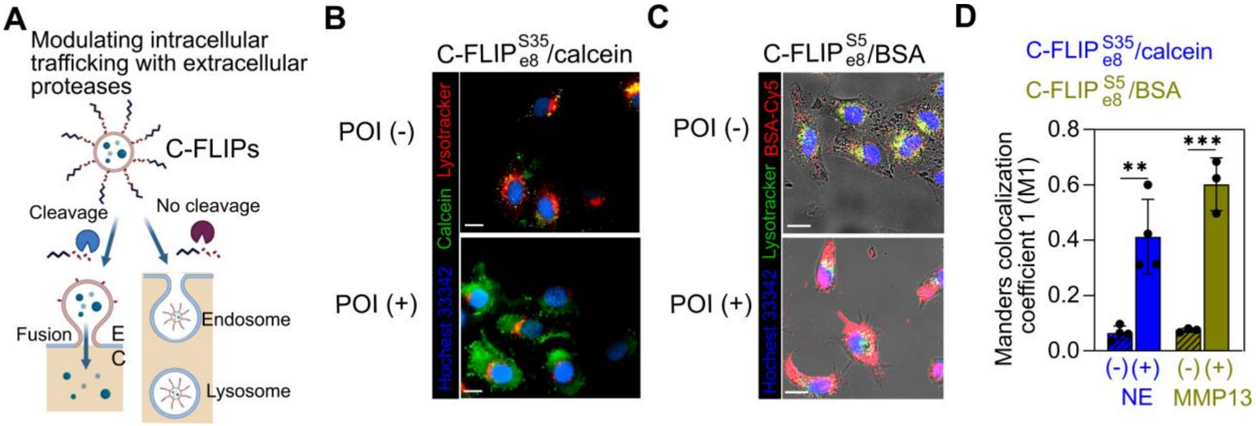


**Figure S3. Cytosolic delivery of representative small molecules and proteins with C-FLIPs. (A)** Cytosolic delivery via C-FLIPs upon protease-activatable membrane fusion. E = extracellular space and C = cytoplasm. Subcellular trafficking of a representative **(B)** small molecule - calcein (green) and **(C)** a model protein - Cy5-labeled bovine serum albumin (BSA; red) via a C-FLIP with a fusion mask e8 conjugated via a NE-cleavable substrate (C-$FLIP^{S35}_{e8}$) or MMP-cleavable substrate (C-$FLIP^{S5}_{e8}$). (-) and (+) indicate the absence and presence of proteases of interest (POI), respectively, which is NE or MMP13. Scale bar = 5µm.The BSA encapsulation rate was 25.7%. **(D)** Manders' colocalization coefficient (M1) that characterizes the fraction of delivered cargos (i.e., calcein or BSA) overlapping with endolysosomes in (B) and (C). *p<0.05 and ****p<0.0001 by Student's t test with respect to the C-FLIPs/cargo without protease pre-cleavage (-).

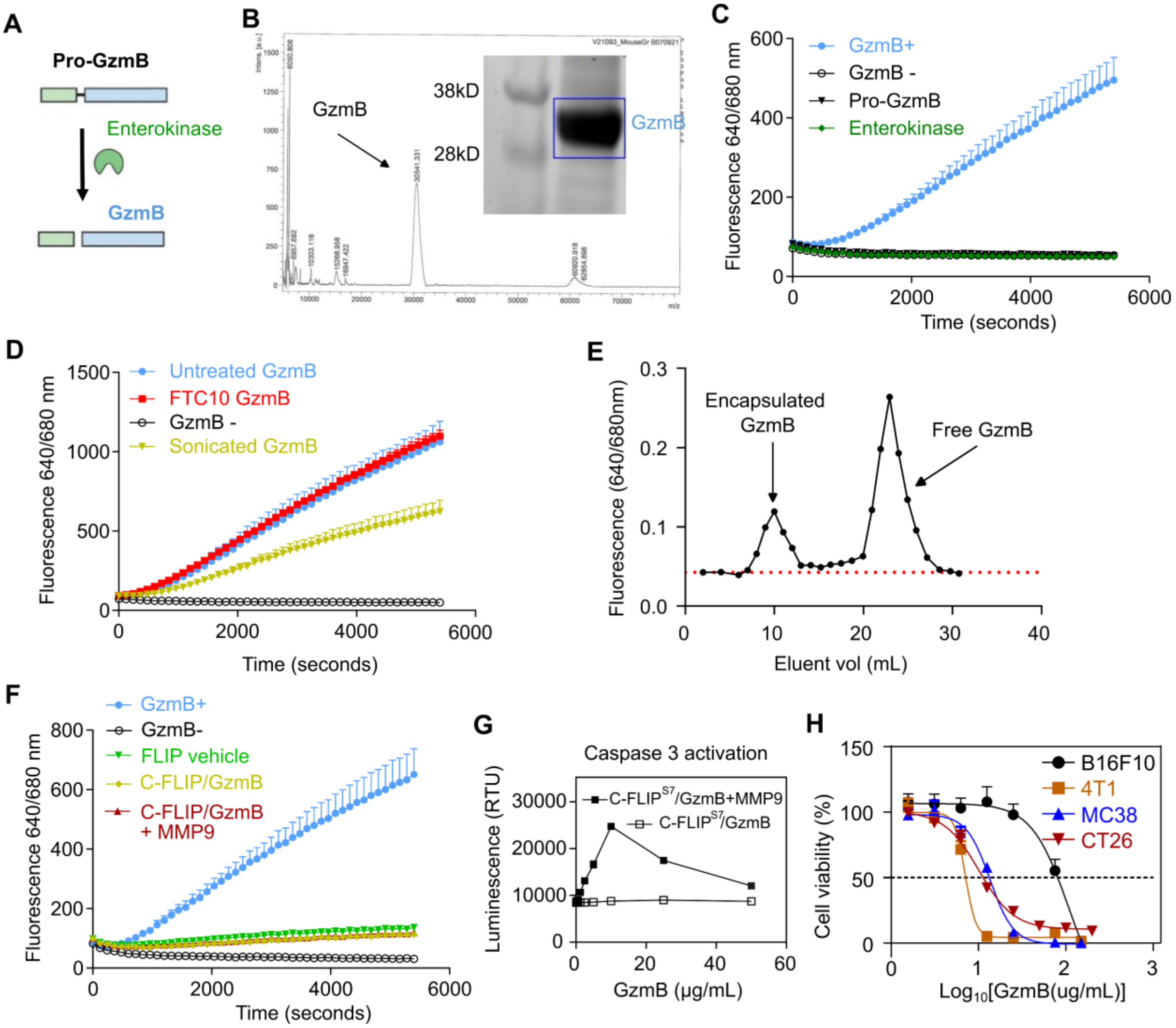


**Figure S4. Synthesis and characterization of C-FLIP/GzmB. (A)** Recombinant pro-granzyme B (Pro-GzmB) is expressed in embryonic FS293 cells (Thermo Fisher Scientific, R79007) and is activated by enterokinase from bovine intestine (Millipore Sigma E5144). **(B)** Molecular weight of GzmB is approximately 30kD measured by MALDI-TOF MS. **(C)** Cleavage of FRET-paired peptide substrate GIEKFD confirms the catalytic activity of the activated GzmB. Fluorescence is activated when fluorophore and its quencher are separated, and the cleavage kinetics was recorded over 1.5 h as indicated. **(D)** GzmB activity remains unchanged after 10 freeze (dry ice)/thaw (37 ºC) cycles (FTC = 10), whereas mild sonication in a bath sonicator reduces the activity of GzmB by nearly 30%. **(E)** Gel filtration to remove unencapsulated Cy5-labeled GzmB from C-FLIP/GzmB. Approximately, 22.6% of GzmB was loaded into the C-FLIP via repeated freezing/thaw processes. The batch-to-batch encapsulation rates vary from 21-27%. **(F)** Catalytic activity of encapsulated GzmB can be temporarily masked by the liposomes since no cleavage of FRET-paired peptide substrate GIEKFD was observed. Pre-activated C-FLIP/GzmB = C-FLIP/GzmB + recombinant MMP9 for 2 h incubation before incubating with FRET-paired GIEKFD. The MMP9 was not removed as it cannot cleave the GIEKFD. Non-activated C-FLIP/GzmB was not incubated with the recombinant MMP9 beforehand. **(G)** Luminescent assay demonstrates the presence of active caspase 3 in the lysed KP cells that were pre-treated with pre-cleaved C-FLIP/GzmB. **(H)** Pre-activated TC-$FLIP_{e8}^{Q7}$/GzmB demonstrates dose-dependent cytotoxicity against a variety of mouse cancer cell lines

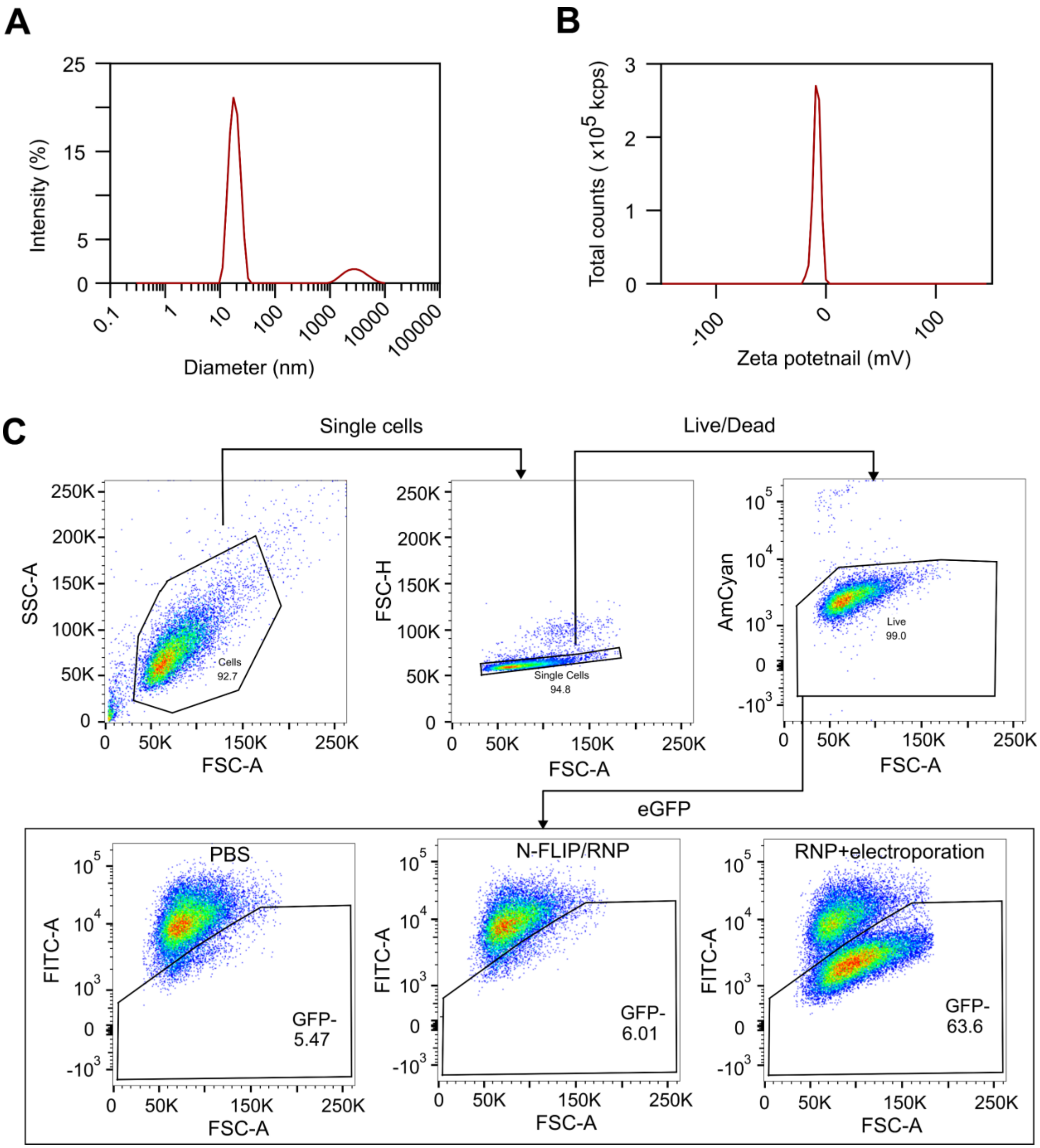


**Figure S5. Gating of eGFP knockout from HeLa d2eGFP cells by ribonucleoprotein particles (RNPs).** The RNPs of CRISPR-Cas9 enzyme and guide RNA was delivered via MMP9-cleavable C-FLIP[S7] or control formulations. **(A)** Hydrodynamic size and **(B)** surface charge of RNPs with Cas9 enzyme and guide RNA. **(C)** Single cell suspension was gated for cells -> single cells -> live/dead cells -> and GFP (FITC channel) negative populations. It is noted that the dot plots of PBS, RNP, RNP via electroporation and pre-cleaved C-FLIP/RNP have been displayed in **Fig.3H**.

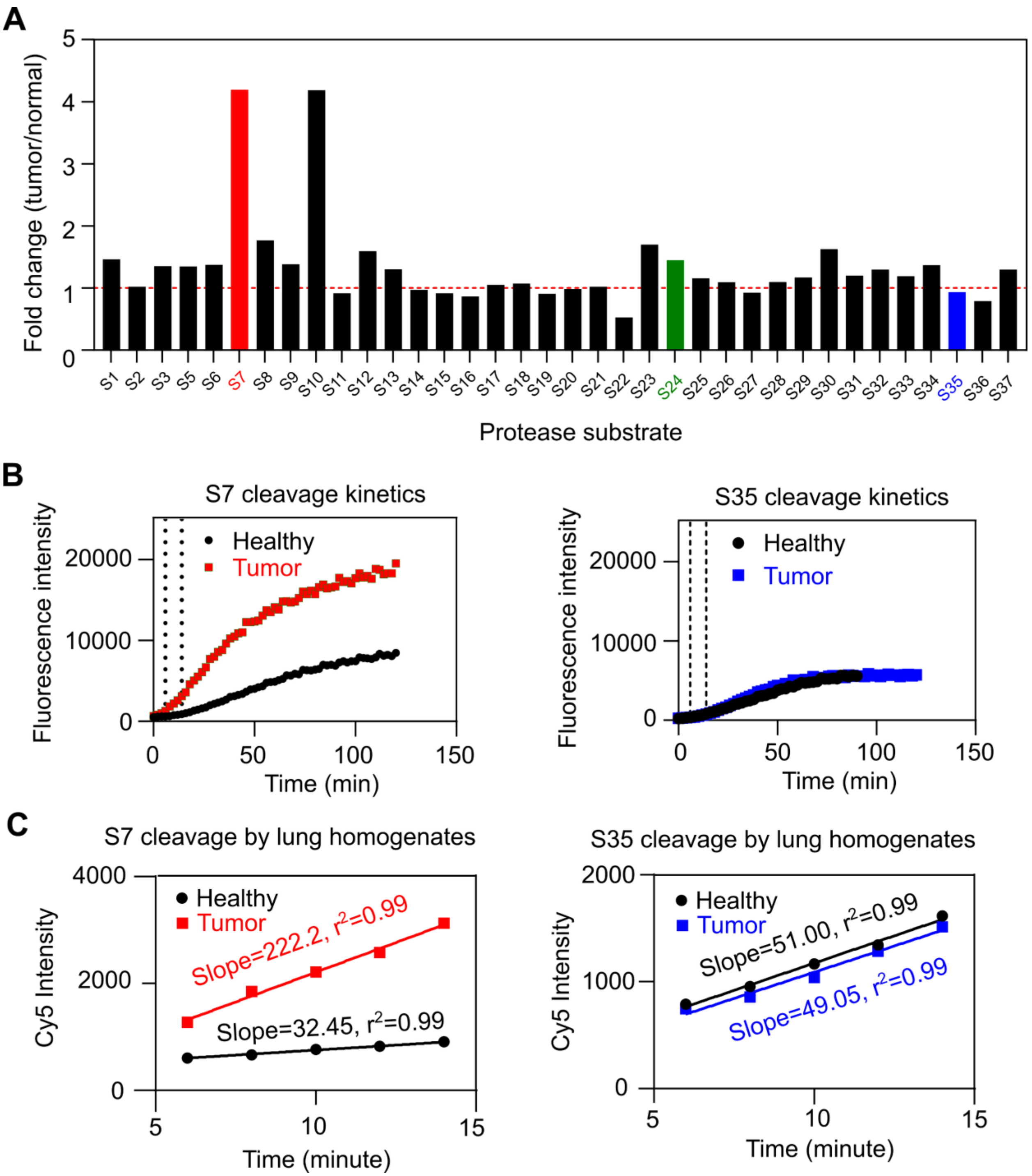


**Figure S6. Protease dysregulation in KP lung tumors**. **(A)** Screening of a library of peptide substrates cleaved by tumor-bearing lung homogenates. The detailed sequences of peptides are referred to **Table S1**. Half a million of KP cells were intravenously injected to wild-type C57BL/6 mice via tail veins. The lungs with tumor nodules were harvested 2 weeks post tumor inoculation. The lung tissues were gently homogenized using a GentleMACS Tissue Dissociator with C tubes. A panel of 36 Förster resonance energy transfer (FRET)–paired peptides were incubated with the tissue homogenates, and fluorescence activation was recorded for 120 mins. Initial cleavage rate of each substrate was calculated [refer to (C)]. The fold change was calculated by normalizing the rate of the tumor group to that of the healthy group. (**B)** Representative cleavage kinetic profile of peptide substate S7 and S35 by the tissue homogenates from healthy lungs and tumor-bearing lungs, respectively. **(C)** Initial cleavage rate of S7 and S35 by KP tumor-bearing lung homogenates during an 8-minute period (6 to 14 mins after recombinant protease and peptide substrate incubation*)*.

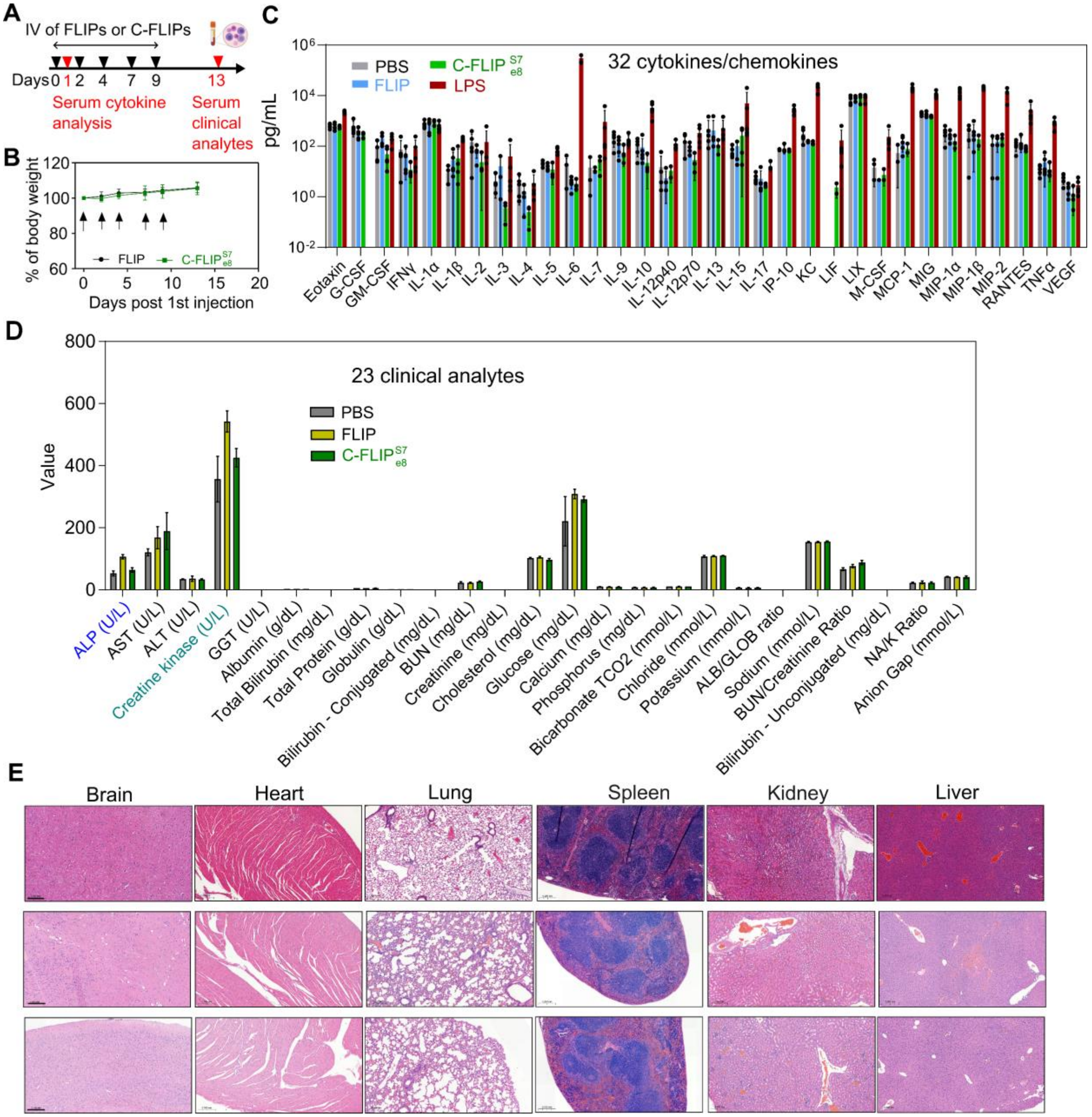


**Figure S7. Toxicity evaluation of intravenously injected C-FLIP. (A)** Experimental timeline for assessing *in vivo* toxicity of 5 doses of the FLIP and MMP9-activatable C-FLIP vehicle with a fusion mask e8 (C-$FLIP_{e8}^{S7}$) in wild-type C57BL/6 mice. Liposomes were intravenously injected at indicated times (black arrows) at a dose of 30 mg/kg. Sera were sampled on Day 1 (i.e., 24 h after the 1st dose) for serum cytokine analysis and on Day 11 (i.e., 4 days after the last dose) for general clinical chemistry. **(B)** Monitoring of body weight over 5 injections of the FLIP and C-FLIP vehicle. **(C)** Assessing 32 cytokines and chemokines in serum samples collected in the wild-type C57BL/6 mice treated with the FLIP and C-$FLIP_{e8}^{S7}$. The serum samples were collected 24h post intravenous injection of the first dose for cytokine analysis by Eve Technologies (Calgary, CAN) using a Luminex-based multiplex assay. G-CSF=granulocyte colony-stimulating factor, GM-CSF=granulocyte-macrophage colony-stimulating factor, IFN-γ=interferon-gamma, IL-1α=interleukin-1 alpha, IL-1β=interleukin-1 beta, IL-2=interleukin-2, IL-3=interleukin-3, IL-4=interleukin-4, IL-5=interleukin-5, IL-6=interleukin-6, IL-7=interleukin-7, IL-9=interleukin-9, IL-10=interleukin-10, IL-12p40=interleukin-12 subunit p40, IL-12p70=interleukin-12 subunit p70, IL-13=interleukin-13, IL-15=interleukin-15, IL-

17=interleukin-17, IP-10=interferon gamma-induced protein-10, KC=keratinocyte chemoattractant, LIF=leukemia inhibitory factor, LIX=lipopolysaccharide-induced CXC chemokine, MCP-1=monocyte chemoattractant protein-1, M-CSF=macrophage colony-stimulating factor, MIG=monokine induced by interferon gamma, MIP-1α=macrophage inflammatory protein-1 alpha, MIP-1β=macrophage inflammatory protein-1 beta, MIP-2=macrophage inflammatory protein-2, RANTES=Regulated on activation, normal T cell expressed and secreted, TNFα=tumor necrosis factor-alpha, and VEGF-A=vascular endothelial growth factor-A. GM-CSF and IL-3 are significantly higher in the cationic FLIP group than PBS-treated counterparts, while the C-$FLIP_{e8}^{S7}$ results in the similar level of GM-CSF and IL-3 to that of the PBS group. **(D)** A panel of 25 serum analytes in the mice injected with 5 doses of the FLIP and C-$FLIP_{e8}^{S7}$ vehicle. Two analytes (inset), alkaline phosphatase (ALP) and creatinine kinase (CK), are significantly higher in the cationic FLIP group than PBS- and C-$FLIP_{e8}^{S7}$-treated groups, suggesting that blocking FLIP with negatively charged peptides lowers the toxicity to the liver and kidney. ALP=alkaline phosphatase, AST=aspartate transaminase, ALT=alanine transaminase, GGT=gamma-glutamyl transferase, BUN=blood urea nitrogen, TCO2=total carbon dioxide, ALB/GLOB=albumin/globulin, and NA/K=sodium/potassium. **(E)** H&E staining of healthy mice dosed with PBS (top row), FLIP (middle row), and C-FLIP (bottom row) for 5 doses. Liposomes were injected at a 30 mg/kg per dose. The scale bars = 200 µm.

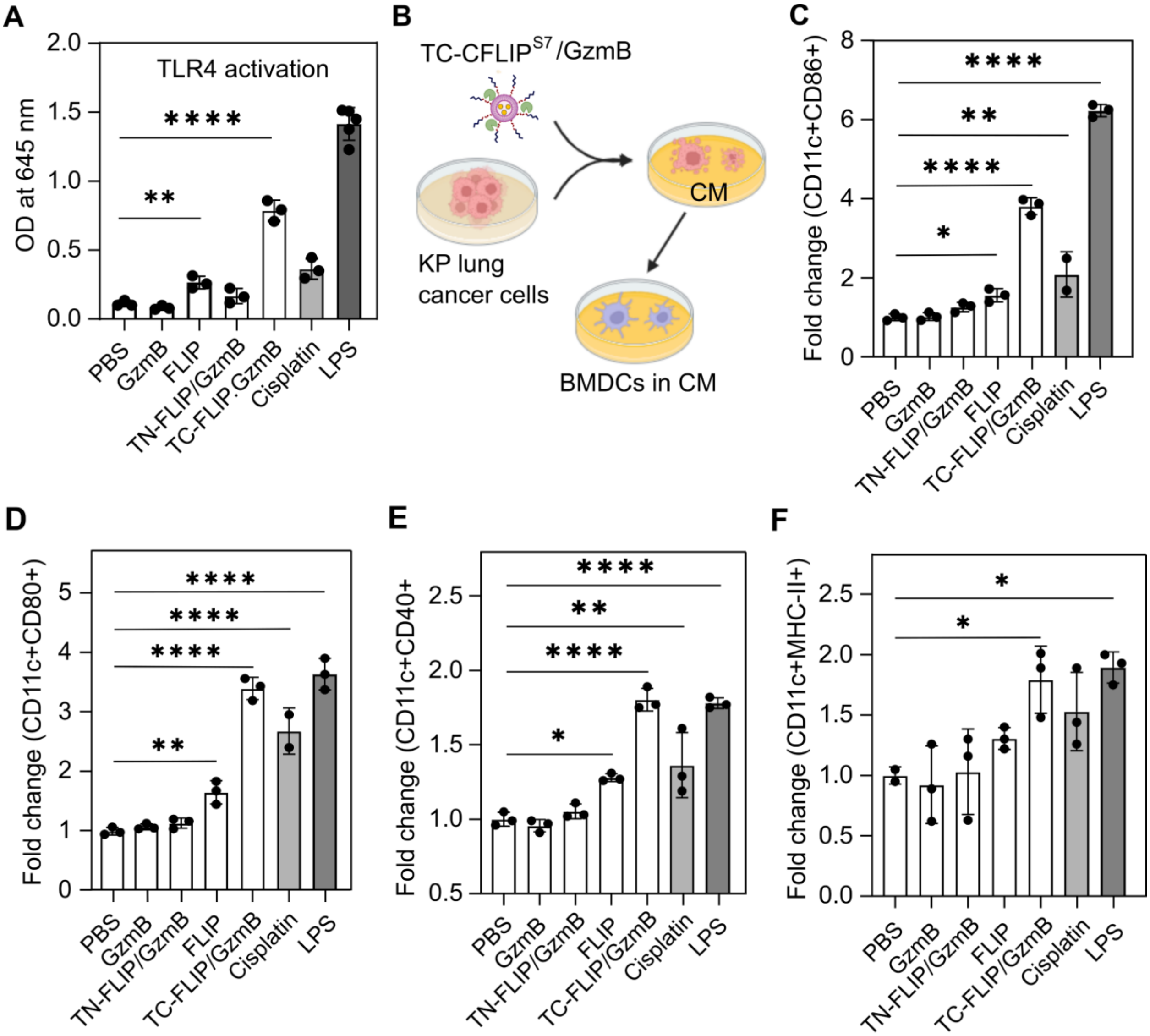


**Figure S8. In vitro activation of dendritic cells by pre-activated TC-FLIP/GzmB**. **(A)** Condition*ed* media (CM) of KP cells treated with FLIP, TC-$\text{FLIP}_{e8}^{S7}$/GzmB, cisplatin and LPS (two positive controls) triggers TLR4 signaling pathway validated using a HEK-TLR4 reporter cell line. HMGB1 is reported to activate the TLR4 pathway. **(B)** Scheme of activation of bone marrow-derived dendritic cells (BMDCs) by CM treated with TC-$\text{FLIP}_{e8}^{S7}$/GzmB. The BMDCs were then examined for various activation biomarkers. Upregulation of **(C)** CD86+, **(D)** CD80+, **(E)** CD40+, and **(F)** MHC-II+ in the BMDCs (CD11c+) after 24h incubation with CM isolated from the KP cells treated with different formulations. The expression of all activation markers was normalized to their quiescent state of untreated BMDCs. Statistical analysis was performed using one-way ANOVA with Dunnett's multiple comparisons with respect to PBS group, with *p<0.05, **p<0.01, and ****p<0.0001. All error bars represent standard deviations.

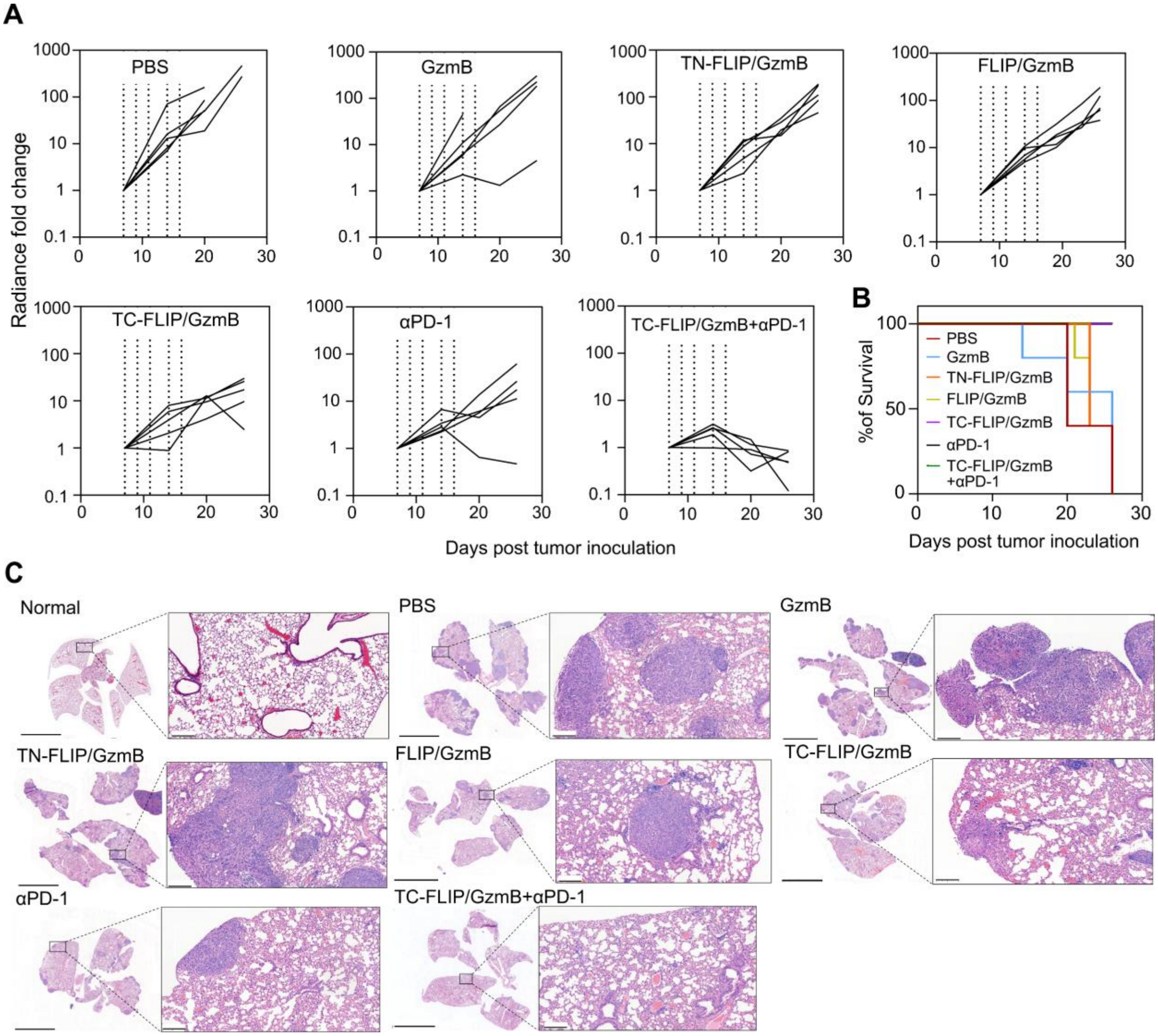


**Figure S9. TC-FLIP/GzmB alone significantly suppresses tumor growth and further synergizes with immune checkpoint blockade in a KP lung metastasis model. (A)** *In vivo* monitoring of tumor proliferation in a KP mouse lung model with *in vivo* imaging system for luciferin luminescence. Wild-type C57Bl/6 mice receiving half a million luciferase-expressing KP cells were treated by PBS, GzmB, targeted non-cleavable fusogenic liposomes/GzmB (TN-FLIP/GzmB), fusogenic liposomes/GzmB (FLIP/GzmB), targeted conditional fusogenic liposomes/GzmB (TC-FLIP/GzmB), programmed death receptor 1 antibody (αPD-1), and combination of TC-FLIP/GzmB and αPD1 (TC-FLIP/GzmB+αPD-1) for 5 doses at a 3 mg/kg of GzmB, and/or 10 mg/kg of PD-1 antibody. The 2$^{nd}$ and 3$^{rd}$ mice (white arrow) in the TC-FLIP/GzmB group were accidently exchanged for positions since Day 14. The tumor burden fold change was calculated accordingly. Dotted vertical lines indicates the dosing of regimens. **(B)** Kaplan survival curve of lung tumor-bearing mice treated with different regimens. ***p*<0.05 and **p*<001 by Long-rank (Mantel-Cox) test with respect to PBS group, **(C)** Hematoxylin and Eosin staining reveals reduced tumor burdens in the TC-FLIP/GzmB alone and TC-FLIP/GzmB+αPD-1 groups. Tumor nodules were visualized in H&E staining of mouse lungs. The lungs were harvested at terminal points (according to euthanasia criteria) or 2 days post last treatment, whichever came first. Scale bar=5 mm and 0.2 mm in the insets.

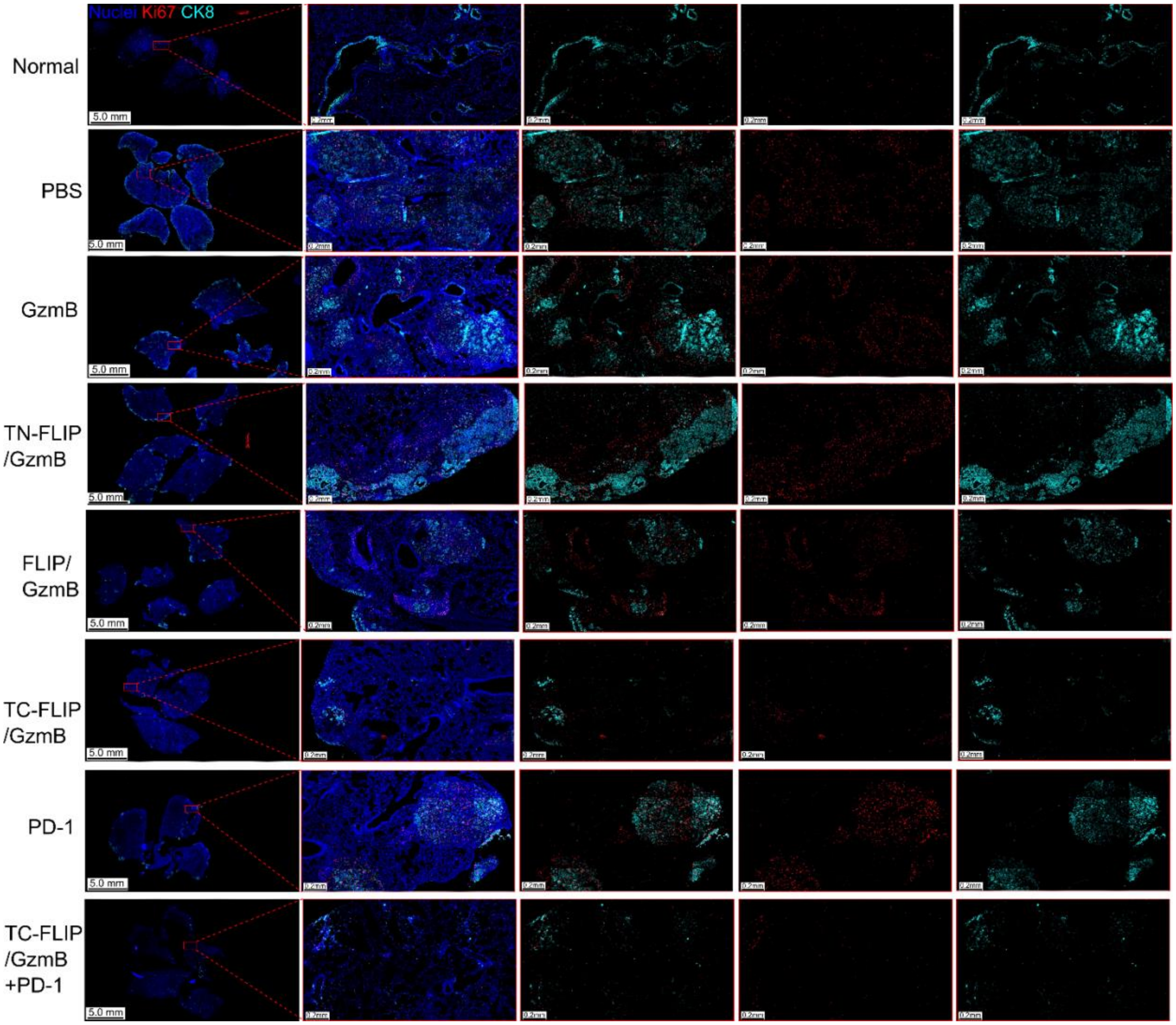


**Figure S10. TC-FLIP/GzmB alone or TC-FLIP/GzmB with PD-1 blocking antibodies downregulate cell proliferation biomarker - Ki67.** Formalin-fixed lung tissue were stained respectively for nuclei (blue), Ki67 (red) - a cell proliferation biomarker, cytokeratin 8 (CK8; Cyan) – a cytoskeletal marker of epithelial cells (cancer cells). Lung tumor (KP)-bearing mice were treated with PBS, granzyme B (GzmB), targeting/non-cleavable fusogenic liposomes/GzmB (TN-FLIP/GzmB), fusogenic liposomes/GzmB (FLIP/GzmB), targeted conditional fusogenic liposomes/GzmB (TC-FLIP/GzmB), programmed cell death protein 1 blocking antibody (αPD-1), and combination of TC-FLIP/GzmB and αPD-1 (TC-FLIP/GzmB+αPD-1 for 5 doses at a 3 mg/kg of GzmB or a 10 mg/kg of αPD-1. The lungs were harvested at terminal point (according to euthanasia criteria) or 2 days last treatment, whichever came first. Scale bar=5 mm or 0.2 mm in magnified images.

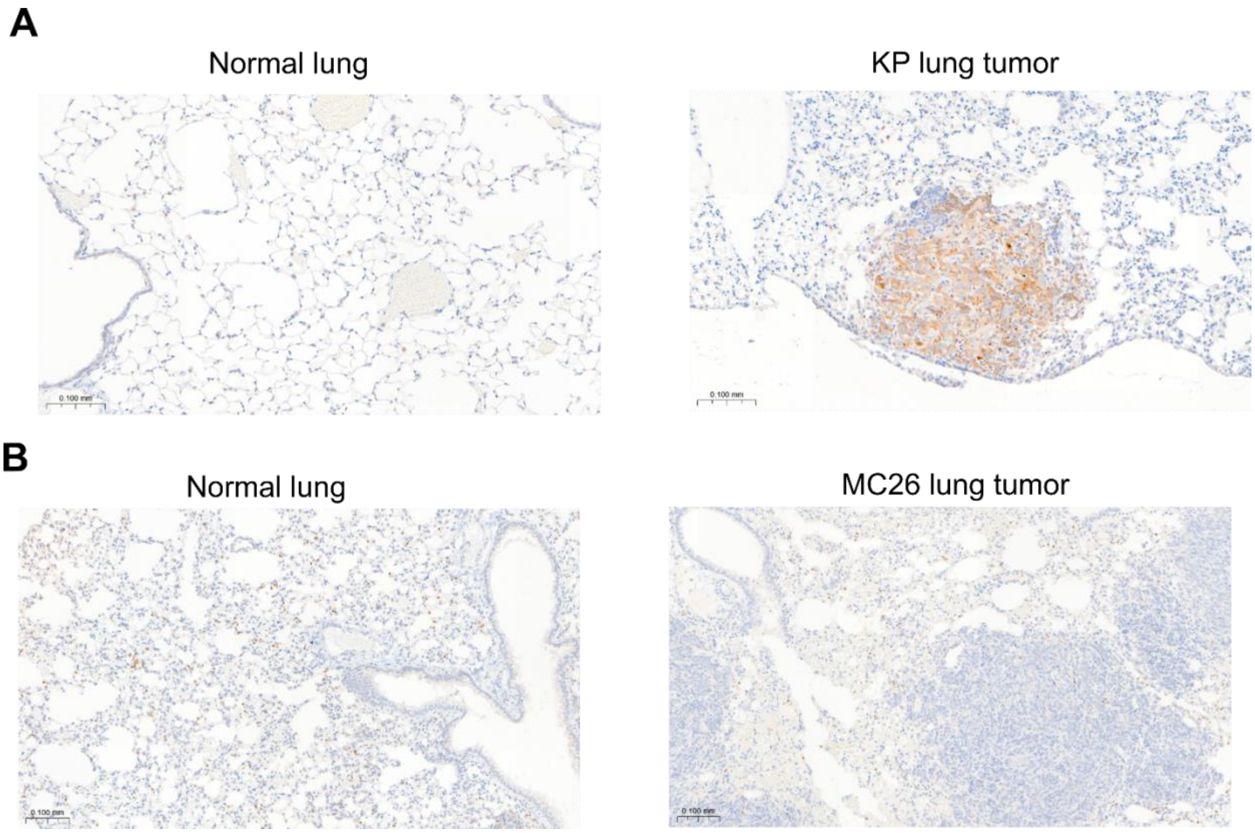


**Figure S11. Immunohistochemical staining of gasdermin E expression in healthy lungs and lung metastases.** Gasdermin E staining in **(A)** healthy lungs and KP lung tumors in the C57BL/6 mice, and in **(B)** healthy lungs and CT26 lung metastasis in the BALB/c mice. Gasdermin E is highly up-regulated in the nodules of KP lung tumors, but barely expressed in the nodules of CT26 lung metastasis. The lung tissues were harvested 14 days post tumor cell inoculation and fixed with 4% PFA. Scale bar = 100 µm

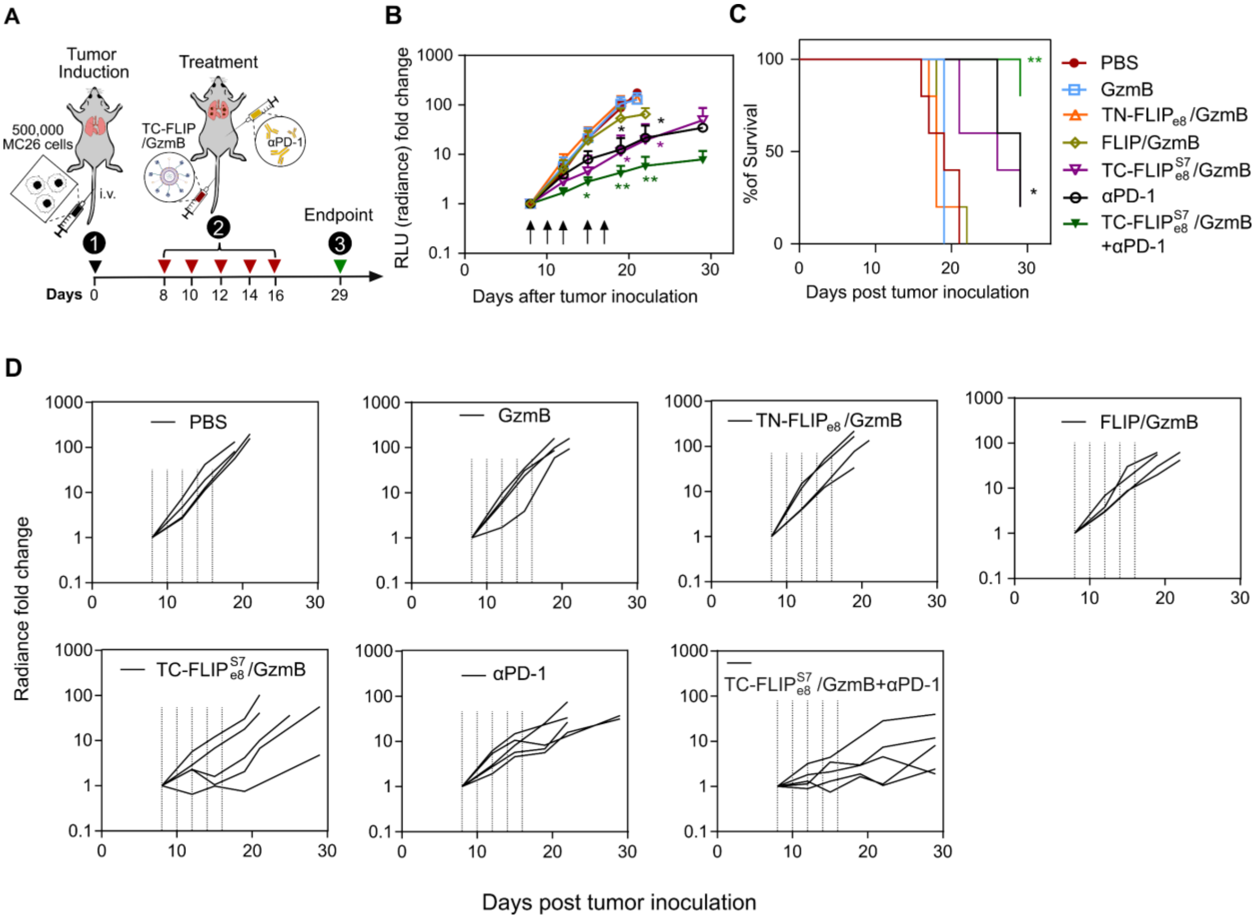


**Figure S12**. **TC-FLIP/GzmB alone suppresses tumor growth and therapeutically synergizes with immune checkpoint blockade in a tumor model of CT26 lung metastasis. (A)** Timeline for therapeutic efficacy study in a CT26 lung transplant tumor model. Half a million of luciferase-transfected CT26 colon cancer cells were injected intravenously into wild-type C57BL/6 mice via tail veins (n=5 per group). Treatments were administered on Day 7, 9, 11, 14 and 16 (pointed by red arrows) for 5 doses in total and tumor burdens were regularly monitored via the IVIS imaging. **(B)** Longitudinal monitoring of tumor burdens and **(C)** Kaplan survival curves of the CT26 lung metastasis-bearing mice treated with different regimens. The mice were treated with i) i.v. PBS, ii) i.v. GzmB 3 mg/kg per dose, iii) i.v. TN-FLIP/GzmB (non-cleavable FLIP with GzmB encapsulated) at 3 mg/kg GzmB, iv) i.v. MMP9-activatable TC-FLIP/GzmB at 3 mg/kg GzmB, v) i.p. αPD-1 (immune checkpoint PD-1 antibody) at 10mg/kg per dose, and vi) i.v. TC-FLIP/GzmB (3mg/kg per dose) in combination with i.p. αPD-1 (10 mg/kg per dose). *in vivo* monitoring of tumor proliferation in a CT26 lung metastasis mouse model. **$p$<0.05 and *$p$<001 by Long-rank (Mantel-Cox) test with respect to PBS group. **(D)** CT26 lung metastasis proliferation profiles of every treated individual mouse.

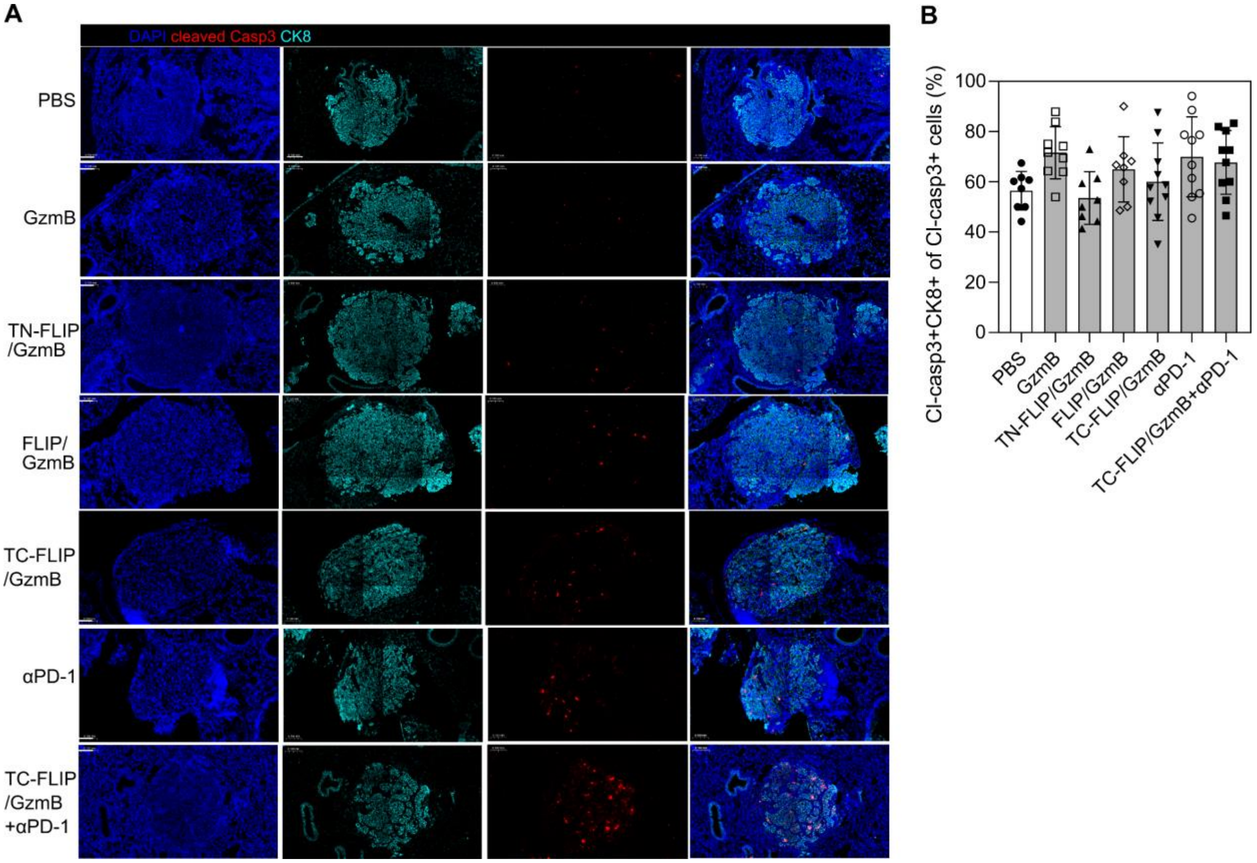


**Figure S13**. **TC-FLIP/GzmB alone or with αPD-1 results in higher cleavage of caspase 3 in lung tumors. (A)** Representative immunofluorescent staining against cleaved caspase 3 (red), abbreviated as cl-casp3, in the lungs and tumor nodules of the mice treated with indicated regimens. The mice were euthanized 24 h after the second dose. DAPI stain nuclei. CK8 (cytokeratin 8) is highly expressed in non-small-cell lung cancer cells. **(B)** Fraction of Cl-casp3+CK8+ cells in the cells undergoing programmed cell death (cl-casp3+) in the tumor nodules. Approximately 50-90% dying cells belongs to the cancer cell population in all treated group without statistical significance. Statistical analysis was performed using one-way ANOVA with Dunnett's multiple comparisons with respect to the PBS group. Each open or closed symbol represents one mouse. The group size n= 8 to 10 mice. All error bars represent standard deviations (SDs). Each open or closed symbol represents one mouse. The group size n= 8 to 10 mice.

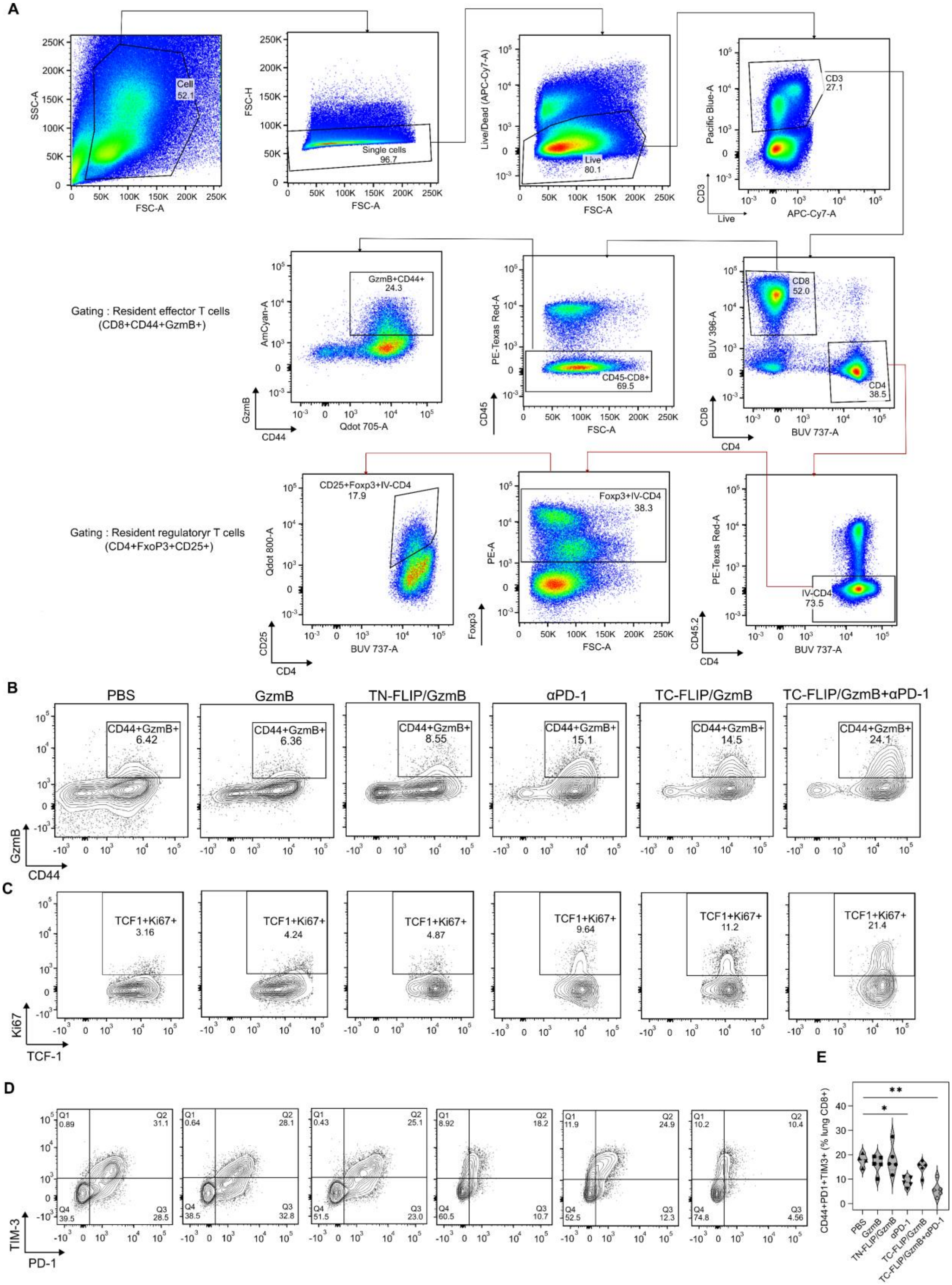
A
Cell
52.1
SSC-A
FSC-A
FSC-H
Single cells
96.7
Live/Dead (APC-Cy7-A)
Live
80.1
Pacific Blue-A
CD3
27.1
APC-Cy7-A
CD3
Live
Gating : Resident effector T cells
(CD8+CD44+GzmB+)
GzmB+CD44+
24.3
AmCyan-A
Qdot 705-A
GzmB
CD44
PE-Texas Red-A
CD45-CD8+
69.5
CD45
BUV 396-A
CD8
52.0
CD4
38.5
BUV 737-A
CD8
CD4
Gating : Resident regulatoryr T cells
(CD4+FxoP3+CD25+)
CD25+Foxp3+IV-CD4
17.9
Qdot 800-A
CD25
CD4
Foxp3+IV-CD4
38.3
PE-A
Foxp3
IV-CD4
73.5
CD45.2
B
PBS
GzmB
TN-FLIP/GzmB
αPD-1
TC-FLIP/GzmB
TC-FLIP/GzmB+αPD-1
CD44+GzmB+
6.42
6.36
8.55
15.1
14.5
24.1
GzmB
CD44
C
TCF1+Ki67+
3.16
4.24
4.87
9.64
11.2
21.4
Ki67
TCF-1
D
Q1 0.89
Q2 31.1
Q4 39.5
Q3 28.5
Q1 0.64
Q2 28.1
Q4 38.5
Q3 32.8
Q1 0.43
Q2 25.1
Q4 51.5
Q3 23.0
Q1 8.92
Q2 18.2
Q4 60.5
Q3 10.7
Q1 11.9
Q2 24.9
Q4 52.5
Q3 12.3
Q1 10.2
Q2 10.4
Q4 74.8
Q3 4.56
TIM-3
PD-1
E
CD44+PD1+TIM3+ (% lung CD8+)
**
*
PBS
GzmB
TN-FLIP/GzmB
αPD-1
TC-FLIP/GzmB
TC-FLIP/GzmB+αPD-1

**Figure S14. Phenotypic profiling of T cells in the lungs with tumors over the treatment. (A)** The single cell suspension was gated for single cells -> live/dead cells -> CD3+ cells -> CD8+ or CD4+ populations. Among the CD8+ cells, we gated for CD45- cells (as CD45 flow conjugated antibodies binds to circulating CD45+ cells, while sparing tissue-resident CD45+ cells) -> CD44+GzmB+ (i.e., effector T cells are CD8+CD44+GzmB+). Among the CD4+ cells, we similarly gated for CD45- cells -> Foxp3+ cells -> CD25+ cells. (i.e., regulatory T cells are CD4+Foxp3+CD25+). **(B)** Representative contour plots of effector CD8+ T cells isolated from the KP tumor bearing mice treated with different regimens. The treatment of TC-FLIP/GzmB results in increased and similar infiltration of effector phenotypes into lung tumors to that by the PD-1 antibody (αPD-1), and the combination of TC-FLIP/GzmB and αPD-1 brings more effector T cells into the tumor microenvironment. **(C)** Representative contour plots of lung resident proliferative stem-like CD8+ T cells isolated from the KP tumor bearing mice treated with different regimens. The cells were gated as for CD44+ cells -> TCF-1+Ki67+ (i.e., proliferative stem-like CD8+ T cells are CD8+CD44+TCF1+Ki67+). The treatment of C-FLIP/GzmB increases CD8+ T cell stemness, and the combination therapy further increase the portion of the progenitor population. **(D)** Representative contour plots of terminally exhausted CD8+ T cells (PD-1+Tim-3+) in the lungs isolated from the KP tumor-bearing mice treated with different regimens. The cells were gated for CD44+ cells -> PD-1+ and PD-1+Tim-3+ (i.e., exhausted T cells are CD8+CD44+PD-1+TIM-3+). **(E)** The treatment of TC-FLIP/GzmB slightly decreases the exhaustion of CD8+ T cells, while the combination therapy of TC-FLIP/GzmB and αPD-1 significantly delays the exhaustion of the CD8+ T cells. The statistical analysis of all groups was performed using one way-ANOVA with Dunnett's comparison with respect to the PBS group.
